# Inhibition of protein translation by the DISC1-Boymaw fusion gene from a Scottish family with major psychiatric disorders

**DOI:** 10.1101/005710

**Authors:** Baohu Ji, Kerin K. Higa, Minjung Kim, Lynn Zhou, Jared W. Young, Mark A. Geyer, Xianjin Zhou

## Abstract

The t(1; 11) translocation appears to be the causal genetic lesion with 70% penetrance for schizophrenia, major depression, and other psychiatric disorders in a Scottish family. Molecular studies identified the disruption of the DISC1 (disrupted-in-schizophrenia 1) gene by chromosome translocation at chromosome 1q42. Our previous studies, however, revealed that the translocation also disrupted another gene, Boymaw (also termed DISC1FP1), on chromosome 11. After translocation, two fusion genes (the DISC1-Boymaw (DB7) and the Boymaw-DISC1 (BD13)) are generated between the DISC1 and Boymaw genes. In the present study, we report that expression of the DB7 fusion gene inhibits both intracellular NADH oxidoreductase activities and protein translation. We generated humanized DISC1-Boymaw mice with gene targeting to examine the *in vivo* functions of the fusion genes. Consistent with the *in vitro* studies on the DB7 fusion gene, protein translation activity is decreased in the hippocampus and in cultured primary neurons from the brains of the humanized mice. Expression of Gad67, Nmdar1, and Psd95 proteins are also reduced. The humanized mice display prolonged and increased responses to the NMDA receptor antagonist, ketamine, on various mouse genetic backgrounds. Abnormal information processing of acoustic startle and depressive-like behaviors are also observed. In addition, the humanized mice display abnormal erythropoiesis, which was reported to associate with depression in humans. Expression of the DB7 fusion gene may reduce protein translation to impair brain functions and thereby contribute to the pathogenesis of major psychiatric disorders.

## Introduction

Over twenty years ago, St Clair et al. reported a large Scottish family with major psychiatric disorders (1). In this family, a balanced chromosome translocation t(1;11) co-segregates with schizophrenia, major depression, and other psychiatric disorders, suggesting the translocation as a causal genetic mutation (2). The DISC1 (disrupted-in-schizophrenia 1) gene was subsequently identified from the breakpoint of the translocation on chromosome 1 (3). Since then, the DISC1 gene has been extensively studied and proposed to play important roles in neuronal cell proliferation, differentiation, migration, maturation, and synaptic functions (4–8). In a comprehensive summary of DISC1 studies, Brandon and Sawa provided a recent review on the potential roles of DISC1 in the pathogenesis of psychiatric disorders (9).

Our previous studies, however, found that the translocation also disrupted another gene, Boymaw (also termed DISC1 fusion partner 1, DISC1FP1), on chromosome 11 in the Scottish family (10). After translocation, the DISC1-Boymaw (DB7) and the Boymaw-DISC1 (BD13) fusion genes are generated. The DB7 fusion protein is insoluble (11). Insoluble DISC1 proteins were reported in the postmortem brains of sporadically collected patients with schizophrenia, major depression, and bipolar disorder (12). These findings prompted us to study the functions of insoluble DB7 fusion protein in hopes of elucidating molecular mechanisms underlying the major psychiatric disorders.

In this study, we investigated the effects of full-length DISC1 (DISC1-FL), truncated DISC1, DISC1-Boymaw (DB7), and Boymaw-DISC1 (BD13) protein expression in different cell lines. We found that expression of the DB7 fusion gene inhibits oxidoreductase activities, rRNA synthesis, and protein translation. Humanized DISC1-Boymaw mice were generated via gene targeting to examine the *in vivo* functions of the fusion genes. We confirmed inhibition of protein translation by the DISC1-Boymaw fusion gene in the humanized mice. Molecular and behavioral characterizations of the humanized mice were also conducted.

## Results

### MTT Reduction and DB7 Fusion Gene Expression

The DISC1 gene is mostly expressed in neuronal cells (13, 14). To examine potential functions of the DB7 fusion gene, we conducted our studies in HEK293T cells that share 90% of expressed genes with human brain (15, 16). The cDNA genes encoding DISC1-FL, truncated DISC1 (1–597 aa), DISC1-Boymaw (DB7), and Boymaw-DISC1 (BD13) are tagged with an HA epitope at their C-terminal (Figure 1A). Full-length DISC1-FL, truncated DISC1, and BD13 fusion proteins were abundantly expressed in contrast to low levels of DB7 fusion proteins exclusively found in the pellet fraction (Figure 1A, overexposure in Figure S1A). Similar expression patterns were also observed in transfected COS-7 cells (Figure S1B and S1C). Since we do not know what exact functions the DB7 protein may serve, we first chose an assay to measure the broad cellular activity of the cells transfected with these genes. MTT (3-(4,5-dimethyliazol-2-yl)-2,5-diphenyltetrazolium bromide) reduction assays were therefore conducted (17–20). Surprisingly, only the DB7 fusion gene expression significantly decreased MTT reduction while MTT reduction did not differ between cells expressing DISC1-FL, truncated DISC1, BD13, and GFP control genes (Figure 1B). Decreased MTT reduction by the DB7 fusion gene expression was also observed in both COS-7 and neuroblastoma SH-SY5Y cells (Figure S1D and S1E). We were perplexed by the low expression of the DB7 fusion proteins because all expression constructs contain the same promoter and transcription terminator. One possible explanation could be that the DB7 mRNA is less stable than the mRNA transcripts of the other three genes. To examine such a possibility, bi-cistronic gene constructs were generated (Figure 1C). If the DB7 mRNA were unstable, we would expect little expression of the fluorescence marker gene downstream of the DB7 open reading frame (ORF). Surprisingly, expression of the fluorescence marker gene was not different between cells expressing any of the four bi-cistronic genes (Figure 1C), indicating no difference in their mRNA abundance. To examine whether low level expression of the DB7 fusion proteins may be caused by rapid protein degradation, we constructed fusion genes in which each gene was fused with a fluorescence timer (FT) E5 (Figure 1D) (21). The fluorescence timer E5 proteins change color from green to red, 9 hr after translation. All fusion genes, except the DB7-FT, produced strong co-localized green and red fluorescence (Figure 1D). However, we could only detect weak green fluorescence in the absence of any red fluorescence in cells expressing the DB7-FT proteins, suggesting possible rapid protein degradation. In further support of this conclusion, addition of proteasome inhibitor N-acetyl-L-leucyl-L-leucyl-L-norleucinal (LLnL) to HEK293T cells transfected with the DB7 fusion gene remarkably increased the level of DB7 protein expression (Figure S1F). To provide direct evidence for rapid protein degradation, we performed a proteomic analysis of cells expressing the DB7 fusion gene using the isobaric tags for relative and absolute quantification (iTRAQ) (Figure S2A, S2B, and S2C). Total proteins from the DISC1-FL and the DB7 over-expression cells were labeled with 114 Da, 115 Da, 116 Da, and 117 Da mass tags, respectively. Hundreds of unique DISC1 peptides were identified in cells expressing the DB7 fusion gene, and their average abundance was about 70% of that in cells expressing the DISC1-FL gene. The abundance of a representative DISC1 peptide is shown between cells expressing full-length DISC1 and the DB7 fusion genes (Figure 1E). We conclude that these abundant DISC1 peptides from cells expressing the DB7 fusion gene are likely peptide fragments generated by rapid protein degradation within the cell.

**Figure 1.**
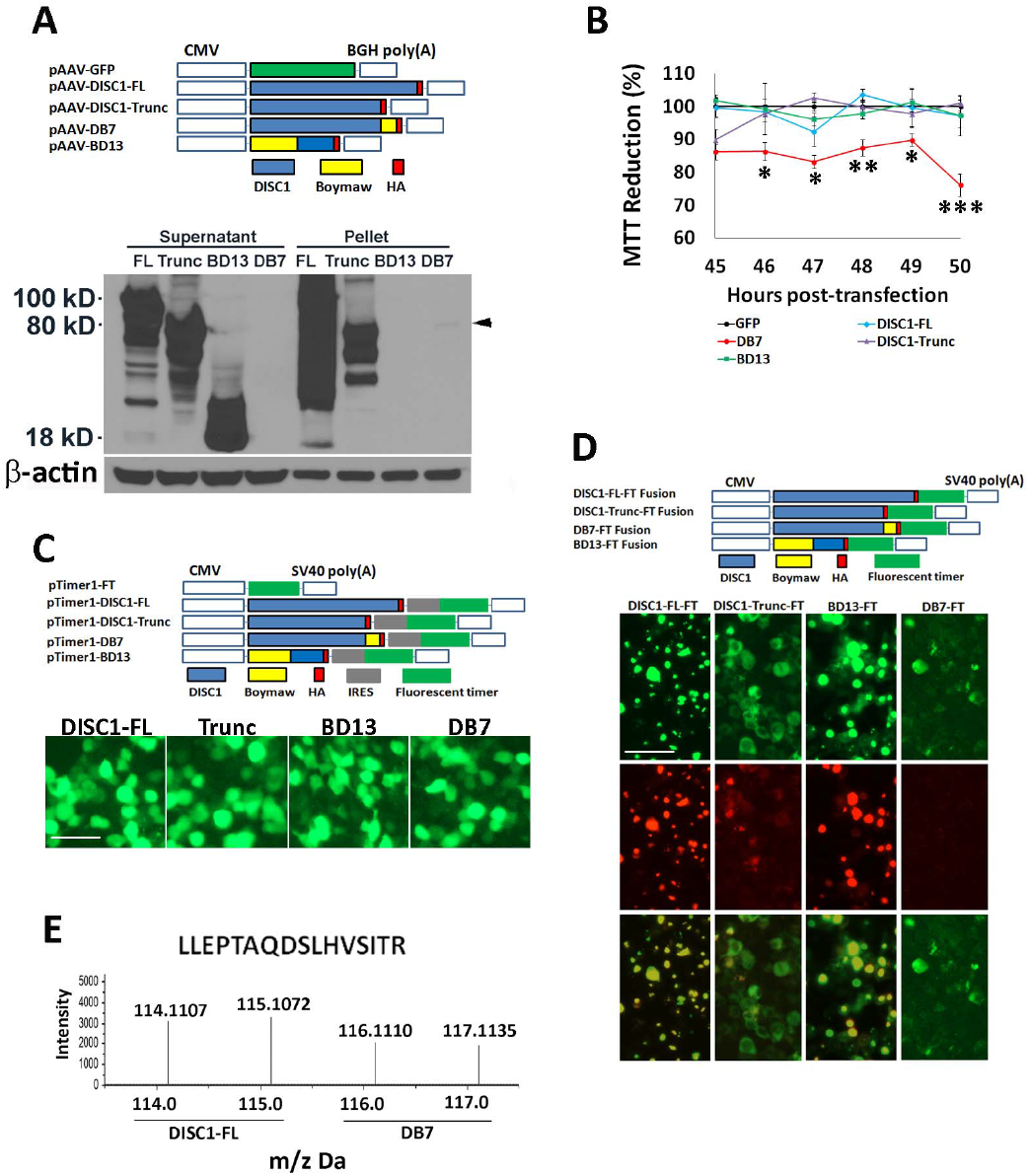
MTT Reduction and Rapid Degradation of DB7 Fusion Protein. **(A)** pAAV plasmid constructs. Each expression construct was tagged with HA epitope. Western blot analyses revealed abundant expression of full-length DISC1 (FL), truncated DISC1 (Trunc), and BD13 (Boymaw-DISC1). The expression of the DB7 (DISC1-Boymaw) was very low (black arrowhead), and restricted to the pellet fraction. Equal amounts of proteins were loaded for the Western blot (see β-actin control). **(B)** MTT reduction assays at multiple time points. There were 4 to 6 replica wells per construct per time point. The mean value of the MTT reduction in cells transfected with GFP was used as the control (100%) to calculate relative MTT reduction for all other constructs at each time point. ANOVA analysis revealed a significant gene effect (F(4,126)=28.09, p < 0.0001). *Post hoc* analyses (Tukey Studentized Range Test) revealed that cells transfected with the DISC1-Boymaw (DB7) construct displayed a significant decrease in MTT reduction at all time points except 45 hr post-transfection. Error bar: SEM. (* p < 0.05, ** p < 0.01, *** p < 0.001). **(C)** bi-cistronic constructs in pTimer1 plasmid vector. Each gene was linked with a fluorescence marker (timer) gene via an IRES site. Green fluorescence was examined 48 hr after transfection of HEK293T cells. There was no difference between any of the four gene constructs. Scale bar: 25 µm. **(D)** The fluorescence timer E5 was in-frame fused with DISC1-FL, truncated DISC1, DB7, and BD13 in pTimer plasmid vector. Expression of the fusion proteins was examined 48 hr after transfection of HEK293T cells. Both green and red fluorescence were readily detected and co-localized in DISC1-FL-FT, DISC1-trunc-FT, and BD13-FT. Weak green, and no red, fluorescence was detected in comparable number of cells expressing DB7-FT only after overexposure. Scale bar: 25 µm**. (E)** In iTRAQ experiments, proteins from cells expressing the full-length DISC1 gene were labeled with 114 Da and 115 Da mass tags; proteins from cells expressing the DB7 gene were labeled with 116 Da and 117 Da mass tags. Abundant human DISC1 peptides were also detected in cells expressing the DB7 gene.

### Characterization of the Boymaw Gene

Since the truncated DISC1 proteins did not decrease MTT reduction, we wondered whether the Boymaw, rather than the DISC1, was responsible for both decreased MTT reduction and rapid protein turnover of the DB7 fusion proteins. Boymaw was therefore in-frame fused at the C-terminal of a randomly selected gene (the fluorescence timer E5 proteins, FT-Boymaw) to follow the same fusion pattern of the DISC1-Boymaw (DB7) gene (Figure 2A). In contrast to strong green and red fluorescence in cells expressing the control construct FT, weak green fluorescence was detected in the absence of red fluorescence in cells expressing the FT-Boymaw fusion gene (Figure 2B). Consistent with the fluorescence images, Western blot analyses detected abundant control FT proteins with little FT-Boymaw fusion proteins (Figure 2C). A significant decrease of MTT reduction was also observed in cells expressing the FT-Boymaw fusion gene in comparison with the controls (Figure 2D). These data indicate that the Boymaw cause decreased MTT reduction as well as rapid protein turnover of both the FT-Boymaw and the DISC1-Boymaw (DB7) fusion proteins. To investigate whether the inhibition of MTT reduction causes rapid protein turnover or vice versa, a series of C-terminal deletions of the FT-Boymaw gene were generated (Figure 2E). The C4-del was as effective as the FT-Boymaw in decreasing MTT reduction (Figure 2F). However, deletion of a short C-terminal sequence in the C4-del dramatically improved protein stability in contrast to barely detectable FT-Boymaw (FT-B) fusion proteins (Figure 2E). The C2-del had a comparable MTT reduction to the controls (FT) (Figure 2F), but was less abundant (Figure 2E). These data indicate that different, although overlapping, regions of the Boymaw gene contribute to the regulation of rapid protein turnover and inhibition of MTT reduction.

**Figure 2.**
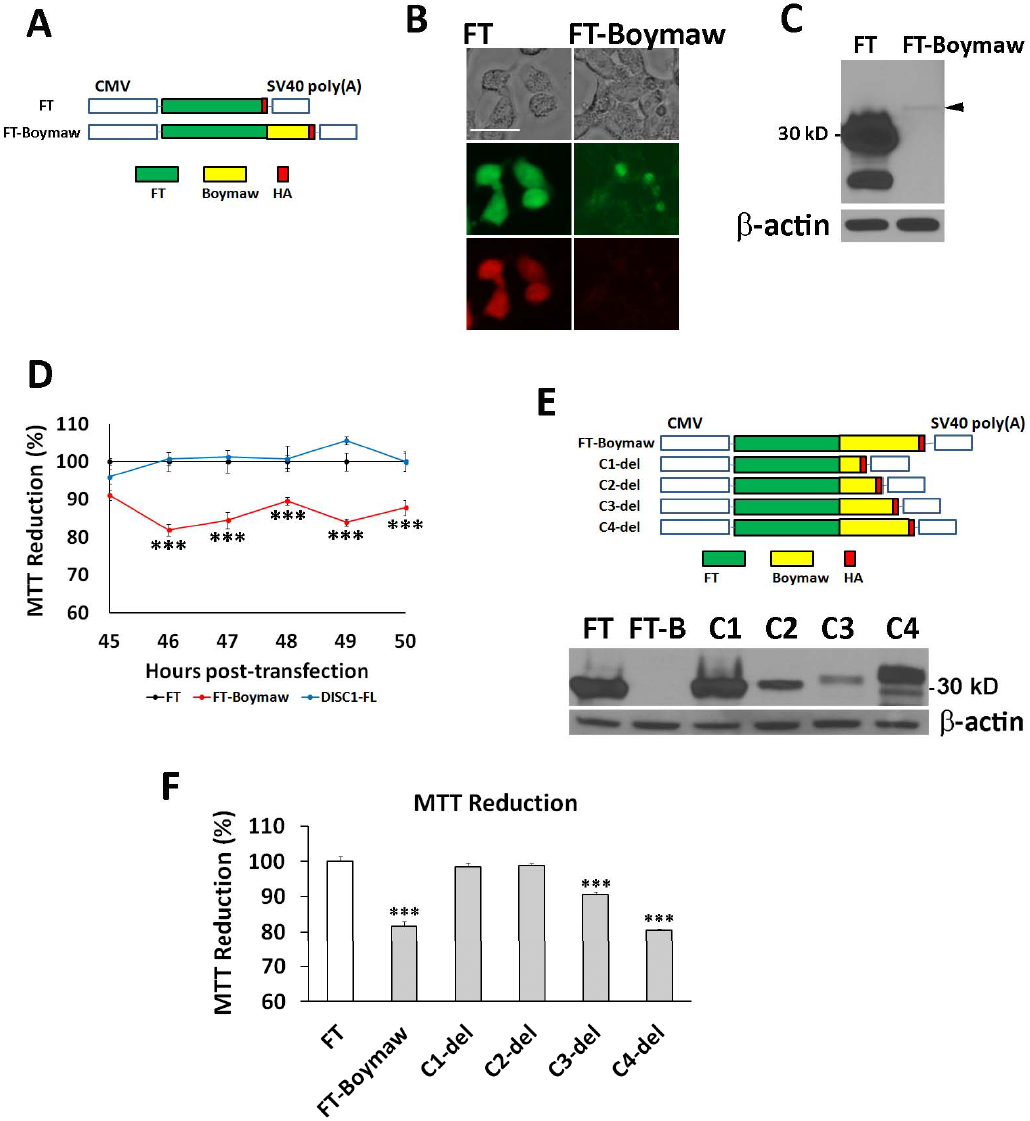
Characterization of the Boymaw Gene. **(A)** Boymaw was fused with a fluorescence timer tagged with HA in pTimer plasmid vector. The pTimer-HA plasmids (FT) were used as controls. **(B)** Expression of the FT-Boymaw fusion proteins was examined 48 hr after transfection of HEK293T cells. While green and red fluorescence of the control fluorescence timer (FT) proteins were very strong, little green and no red fluorescence of FT-Boymaw was detected. Scale bar: 15 µm. **(C)** Western blot analysis barely detected expression of the FT-Boymaw fusion proteins (black arrowhead) in comparison with abundant control FT proteins. **(D)** MTT assays revealed decreased MTT reduction by expression of the FT-Boymaw fusion gene in comparison with the controls and DISC1-FL (F(2,66)=145.12, p < 0.0001). *Post hoc* analyses (Tukey Studentized Range Test) revealed that cells transfected with the FT-Boymaw construct displayed a significant decrease in MTT reduction at all time points except 45 hr post-transfection. There were 4 to 6 replica wells per construct per time point. The mean value of the MTT reduction in cells transfected with FT was used as the control (100%) to calculate relative MTT reduction for the other two constructs at each time point. **(E)** A series of C-terminal deletion of FT-Boymaw fusion gene. Western blot analysis revealed expression of each deletion construct. **(F)** MTT assays were conducted 48 hr after transfection of each deletion construct. There was a significant group effect on MTT reduction (F(8, 27)= 92.14, p < 0.000001). *Post hoc* analyses (Tukey Studentized Range Test) revealed significant MTT reduction in cells transfected with FT-Boymaw, C3-del, and C4-del. There were 4 replica wells per construct. The mean value of the MTT reduction in cells transfected with FT was used as the control (100%) to calculate relative MTT reduction for all other constructs. Error bar: SEM. (* p < 0.05, ** p < 0.01, *** p < 0.001)

### Reduced NADH Oxidoreductase Activities

Our studies found that decreased MTT reduction is not caused by reduced proliferation of the cells expressing the DISC1-Boymaw (DB7) fusion gene (Figure S3A). To rule out potential differences in MTT uptake in living cells, the transfected cells were partially lysed for the MTT assay. DB7 cell lysate display a similar magnitude of MTT reduction deficit to the living cells (Figure 3A). These data therefore indicate that intracellular MTT reduction activities are decreased in DB7 expressing cells. NADH oxidoreductases are major intracellular electron donors in the reduction of MTT (20). We therefore first examined whether NADH concentration was decreased in cells expressing the DB7 gene. No significant difference in intracellular NADH was found between cells expressing DISC1-FL and DB7 genes (Figure S3B). Cytosolic NADH concentration was also examined with NADH fluorescence sensor Peredox (22), and no difference was found either (Figure S3C). To further examine whether NADH oxidoreductase activities were reduced in DB7 expressing cells, recombinant *E coli* NADH alcohol dehydrogenase (ADH), which catalyzes MTT reduction in the presence of NADH, was added to the DISC1-FL and DB7 cell lysates, respectively. Addition of the ADH dramatically increased MTT reduction in the DB7 cell lysate, but not in the DISC1-FL cell lysate (Figure 3B). These data indicate that the DB7 cell lysate have decreased activities of NADH oxidoreductases. To localize the NADH oxidoreductases, we conducted density-gradient ultracentrifugation to isolate both mitochondria and microsomes (fragmented endoplasmic reticulum). Western blot analyses demonstrated enriched expression of cytochrome c in the mitochondrial fraction and cytochrome p450 reductase in the microsomal fraction (Figure 3C). A significant decrease of MTT reduction was observed in both mitochondrial and microsomal fractions from the cells expressing the DB7 gene. Further analysis identified a significant reduction of cytochrome b5 reductase 3 (CYB5R3) proteins in the microsomal fraction of the cells expressing DB7 (Figure 3D, 3E). No differential CYB5R3 expression was detected between DISC1-FL and DB7 in either cell homogenate or mitochondria.

**Figure 3.**
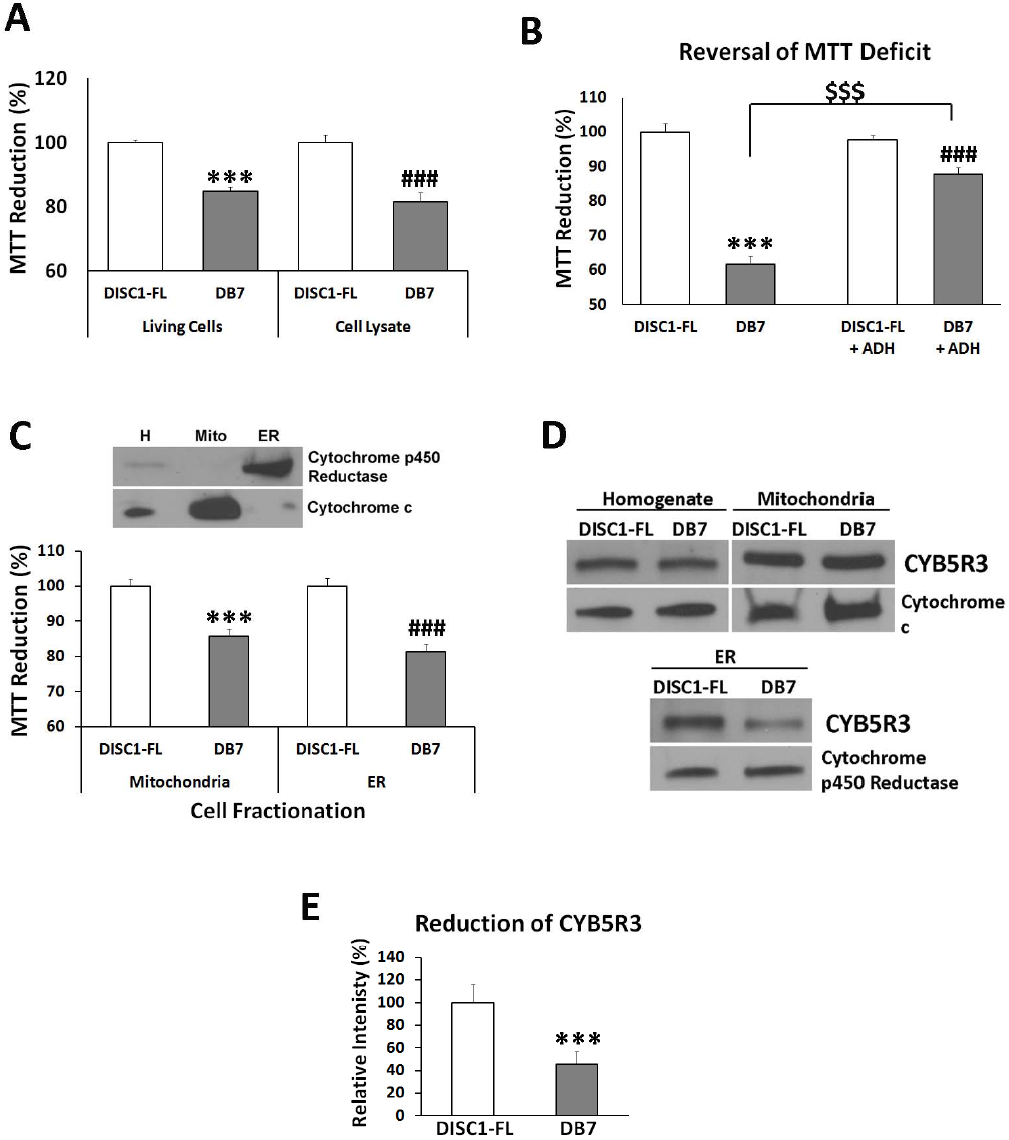
Reduced NADH Oxidoreductase Activities. **(A)** HEK293T cells were transfected with DISC1-FL and DB7 respectively. Two days after transfection, MTT reduction was performed in both living cells and cell lysate respectively. Student’s t-tests (unpaired, two-tailed) were used to compare MTT reduction between DISC1-FL and DB7 in living cells and cell lysate respectively. The mean value of the MTT reduction in cells transfected with DISC1-FL was used as the control (100%) to calculate relative MTT reduction for DB7 in each group. Significantly decreased MTT reduction in DB7 transfection was observed in living cells (n = 3) (t(4)=10.74, p < 0.001).and cell lysate (n = 6) (t(10)=4.92, ^###^ p < 0.001). **(B)** Reversal of MTT reduction deficit in cell lysate. There were 12 replica wells for each gene and treatment. Unpaired student’s t-test (two-tailed) was used for statistical analysis. Significantly decreased MTT reduction was observed in HEK293T cell lysate from DB7 transfection in comparison with DISC1-FL (t(22)=11.50, p < 0.001). *E coli* alcohol dehydrogenase (ADH) can catalyze MTT reduction in the presence of NADH. Addition of the ADH did not completely reverse the deficit of MTT reduction in DB7 lysate in comparison with DISC1-FL lysate (t(22)=4.26, ^###^p < 0.001). However, addition of recombinant *E coli* ADH significantly improved MTT reduction in DB7 cell lysate (t(22)=8.66, ^$$$^ p < 0.001 (comparison between DB7 lysate with and without ADH)). No difference was found in MTT reduction of the DISC1-FL cell lysate with or without ADH. The mean value of the MTT reduction of DISC1-FL samples without ADH was used as the control (100%) to calculate relative MTT reduction for DB7, DISC1-FL+ADH, and DB7+ADH, **(C)** Purification of mitochondria and endoplasmic reticulum (ER) from transfected HEK293T cells using density-gradient ultracentrifugation. Large amount of cells was transfected for each purification experiment. Cytochrome c and cytochrome p450 reductase were used as marker proteins for mitochondria and ER respectively. MTT reduction assays were conducted in purified mitochondria and ER from HEK293T cells transfected with DISC1-FL and DB7 constructs. The value of MTT reduction was further normalized with total amount of proteins in mitochondria and ER, respectively. Significantly reduced MTT reduction was observed in both mitochondria (n = 6)(t(10)=5.53, two-tailed, p < 0.001) and ER (n = 6)(t(10)=6.59, two-tailed, ^###^ p < 0.001) isolated from cells transfected with DB7 expression construct. **(D)** The level of CYB5R3 protein expression was examined with Western blot analysis, and normalized with the expression of ER marker cytochrome p450 reductase (CYPOR). CYB5R3 expression was reduced in the ER of the cells expressing the DB7 fusion gene. No significant difference in CYB5R3 expression was observed between DISC1-FL and DB7 in either cell homogenate or mitochondria after normalization with mitochondrial marker cytochrome c. **(E)** CYB5R3 expression was quantified in ER preparations of each construct (n=4). Unpaired student’s t-test (two-tailed) analysis revealed a significant reduction of CYB5R3 in DB7 (t(6)=18.84, p < 0.001) Error bar: SEM. (* p < 0.05, ** p < 0.01, *** p < 0.001)

To fully characterize the effects of the DB7 fusion gene expression, we conducted proteomic analysis of cells expressing the DB7 fusion gene. We did not find additional candidate NADH oxidoreductases whose expression was significantly decreased (Figure S2A, S2B, S2C, and Table S1). Several reasons could account for the results. First, proteomic analysis can only survey a portion of total cellular proteins. Second, we found that expression of several NADH oxidoreductases was decreased, but did not reach statistical significance because of a limited number of identified peptides. Third, NADH oxidoreductase activities are modulated not only by protein expression but also by protein modifications. The molecular nature of the reduced activities of NADH oxidoreductases in cells expressing the DB7 fusion gene remains to be fully investigated. Among the top 18 differentially expressed proteins ranked according to their p values, however, seven were directly involved in the regulation of protein translation (Figure S2C).

### Inhibition of rRNA Expression and Protein Translation *in vitro*

To investigate whether expression of the DISC1-Boymaw (DB7) fusion gene alters protein translation, we first examined rRNA expression in cells expressing the DISC1-FL, truncated DISC1, DB7, and BD13 genes, respectively. Interestingly, we found a significant reduction of total RNA in cells expressing the DB7 fusion gene (Figure 4A). Reduction of the total RNA was not caused by differential cell proliferation (Figure S3D). Since rRNA makes up more than 80% of total RNA, we quantified the expression of both 28S and 18S rRNAs. Reduction of both 28S and 18S rRNAs was found only in cells expressing the DB7, not the other three genes. Consistent with the effects of the DB7 fusion gene expression, expression of the FT-Boymaw fusion gene also decreased total RNA and rRNA expression (Figure 4B). To provide direct evidence for inhibition of protein translation, we conducted SUnSET (surface sensing of translation) experiments to measure protein translation in the transfected cells (23). In the presence of cycloheximide, an inhibitor of protein translation, we detected little puromycin incorporation in newly synthesized proteins in Western blot analysis (Figure 4C). In pulse-labeling experiments, we found significant reduction of puromycin labeling from cells expressing the DB7 fusion gene, suggesting decreased protein translation activities (Figure 4D, 4E). There was no significant difference in puromycin labeling between cells expressing the other three genes. We confirmed that expression of the FT-Boymaw fusion gene also inhibits protein translation similar to the DB7 fusion gene (Figure 4F, 4G). All together, it appears that it is the Boymaw gene, rather than the DISC1 gene, that contributes to the inhibition of oxidoreductase activities, rRNA expression, and protein translation in both the DISC1-Boymaw and FT-Boymaw fusion genes. We did not observe any changes of these cellular activities in cells expressing the full-length DISC1, truncated DISC1, or the Boymaw-DISC1 fusion genes.

**Figure 4.**
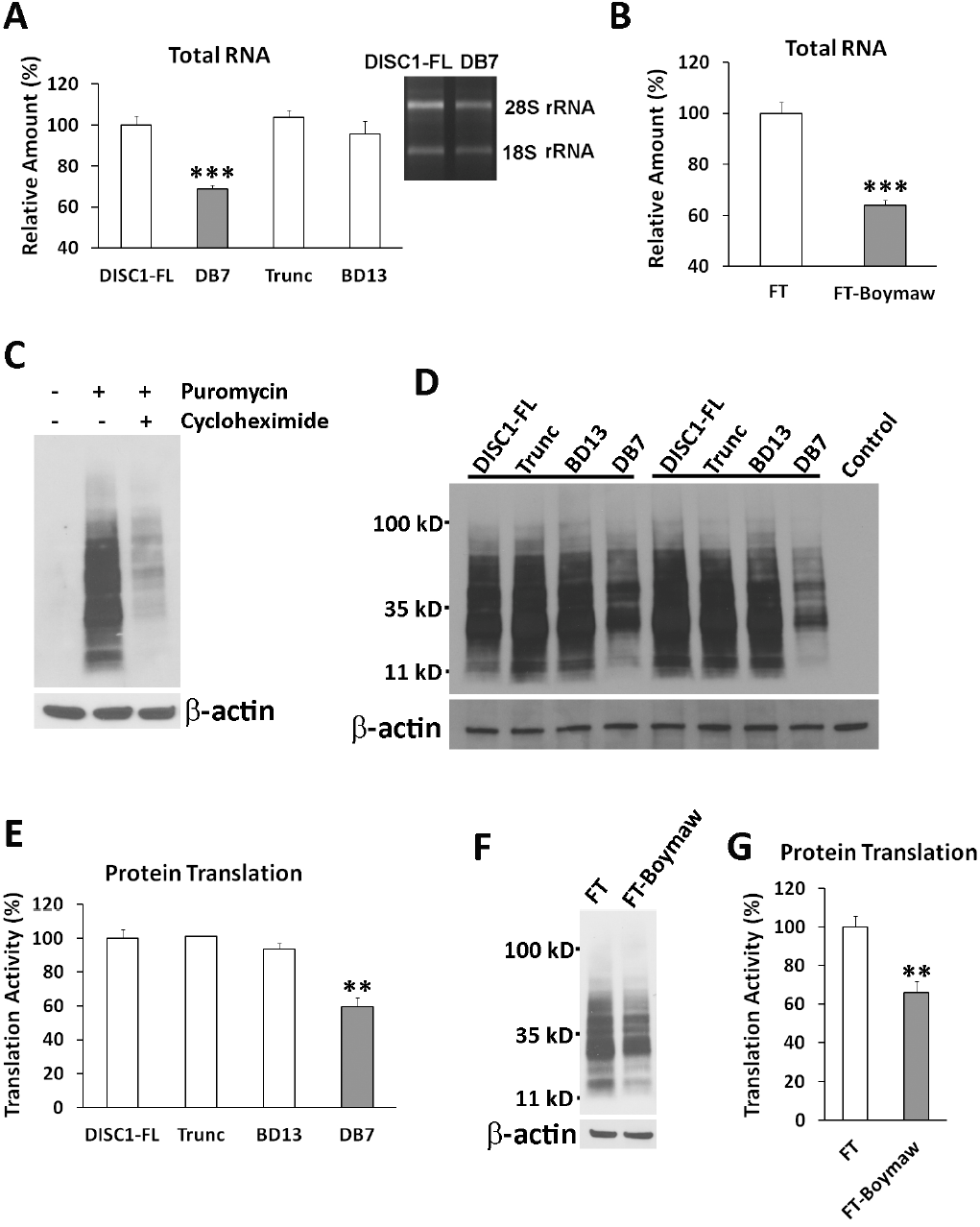
Inhibition of rRNA Expression and Protein Translation. **(A)** Reduction of total RNA in cells expressing the DB7 gene (unpaired, two-tailed student’s t-test, t(10)=9.84, p < 0.001). There were three to six replica wells for each gene construct. Significant reduction of rRNA expression was observed between DB7 and DISC1-FL. The same amount of rRNA reduction was further confirmed in later transfection experiments. Expression of total RNA and rRNA was normalized against the number of cells. The level of total RNA and rRNA expression of the DISC1-FL was used as the reference (100%) to calculate relative expression of the other three groups. **(B)** Reduction of total RNA and rRNA in cells expressing FT-Boymaw fusion gene compared with the control FT gene (unpaired, two-tailed student’s t-test, t(8)=7.6, p < 0.001). There were five replica wells for each construct. Significant reduction of rRNA was observed in cells expressing the FT-Boymaw fusion gene. **(C)** Pulse-labeling of new protein synthesis via incorporation of puromycin. Anti-puromycin antibody has no cross-reactions with HEK293T cellular proteins. A 10-min pulse-labeling generated massive incorporation of puromycin in Western blot analysis. However, the presence of cycloheximide, an inhibitor of protein translation, dramatically reduced the puromycin labeling. **(D)** Inhibition of protein translation. Protein translation activities were measured with SUnSET in cells at 48 hr post-transfection. Remarkable reduction of puromycin labeling was observed in cells expressing the DB7 construct in two representative sets of transfection experiments. **(E)** Quantification of reduction of puromycin incorporation. There were four replica dishes for each construct. There was a significant reduction of protein translation in cells expressing the DB7 construct in comparison with the DISC1-FL construct (unpaired, two-tailed student’s t-test, t(6)=5.03, p < 0.01). The mean value of the incorporation of puromycin in DISC1-FL samples was used as the control (100%) to calculate relative intensity of puromycin incorporation in the other three genes. No difference was found in protein translation activity between the other three constructs. **(F)** There was a reduction of protein translation in cells expressing the FT-Boymaw fusion gene compared with the control FT gene in SUnSET. **(G)** There were six replica dishes for each construct. Significant reduction of puromycin incorporation was observed in cells expressing the FT-Boymaw fusion gene (unpaired, two-tailed student’s t-test, t(10)= 4.48, p < 0.01). Error bar: SEM. (* p < 0.05, ** p < 0.01, *** p < 0.001)

### Generation of Humanized DISC1-Boymaw Mice

To investigate the *in vivo* functions of the DB7 fusion gene, we generated humanized DISC1-Boymaw mice. Human DISC1-Boymaw fusion gene (DB7) and Boymaw-DISC1 fusion gene (BD13) were knocked-in to replace mouse endogenous *disc1* gene (Figure 5A). A bi-cistronic gene, consisting of the two fusion genes connected with an internal ribosome entry site (IRES), was designed to splice with exon 1 of human DISC1 gene. The bi-cistronic cassette was also flanked with two *loxP* sites upstream of a human DISC1 gene for future genetic rescue experiments. Germ-line transmission of the bi-cistronic gene was obtained. To examine whether the bi-cistronic gene is expressed and correctly spliced with exon 1, we conducted RT-PCR analysis of total RNA extracted from the brains of humanized DISC1-Boymaw mice (DISC1^H^) (Figure 5B). A 150 bp DNA fragment was specifically amplified. Further sequencing confirmed the correct splicing (data not shown). Before examination of the expression of the DB7 proteins of the bi-cistronic gene in the humanized mice, we first evaluated eight different anti-DISC1 antibodies using our DISC1 over-expressing cell lysates (Figure S4A). Five antibodies (ab55808, ab62069, ab59017,3D4, and 14F2) with the highest affinities to human DISC1 proteins were selected for Western blot detection of the DB7 fusion protein. Unfortunately, there were extensive non-specific cross-reactions of the antibodies with proteins from mouse brain homogenate in Western blot analysis (data not shown), preventing the detection of the fusion proteins. However, the antibodies have less background in Western blot analyses of mouse primary neuron culture (personal communication with Dr. Hongjun Song). We therefore performed our analyses in cultured primary neurons from neonatal mice. The antibody against the N-terminal residues of human DISC1 protein recognized a 75 kD protein band in Western blot analysis of the primary neurons from the humanized DISC1-Boymaw mice, but not wildtype mice (Figure 5C). Two reasons support this band as the DB7 fusion protein. First, the protein band is about the calculated size of the DB7 fusion protein (73 kD). Second, the absence of this band in wildtype mice rules it out as a non-specific cross-reaction of the antibody. To detect the expression of the BD13 proteins of the bi-cistronic gene, we further screened the DISC1 antibodies in our BD13 over-expressing cell lysate. Two antibodies recognizing the C-terminal residues of human DISC1 proteins displayed some, but not high (compared with anti-HA antibody), affinity to the BD13 proteins (Figure S4B). Due to the low affinity of the two antibodies, we could not detect the expression of the BD13 proteins expected to be around 17 kD (Figure S4C, S4D). To further confirm the expression of the DB7 fusion proteins, immunocytochemical analyses were conducted. Co-localization of the DB7 proteins with neuronal cell marker MAP2 was observed in the primary neurons from the heterozygous DISC1-Boymaw mice (Figure 5D). Expression of the DB7 protein did not cause gross anatomical abnormalities in the brain of the heterozygous DISC1-Boymaw mice (Figure S5). To examine whether expression of the DB7 fusion protein decreases MTT reduction *in vivo*, we performed MTT reduction assays in the brain homogenate of neonatal heterozygous DISC1-Boymaw and wildtype control mice. A significantly decreased MTT reduction was observed in the heterozygous DISC1-Boymaw mice (Figure 5E). Western blot analysis also confirmed a significant decrease of Cyb5r3 proteins in ER isolated from the brains of the heterozygous DISC1-Boymaw mice (Figure 5F).

**Figure 5.**
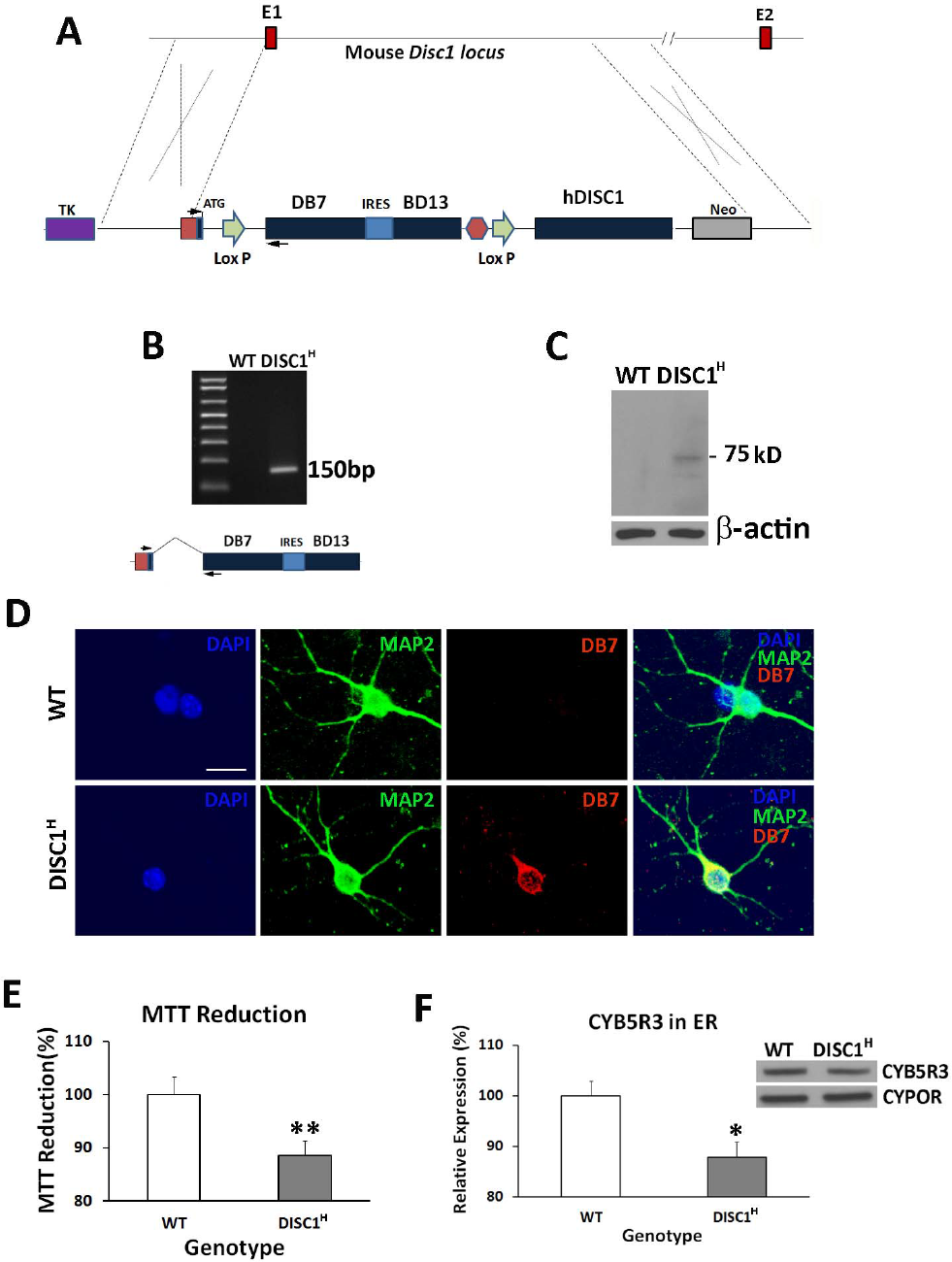
Generation of Humanized DISC1-Boymaw Mice. **(A)** Both the DISC1-Boymaw and the Boymaw-DISC1 fusion genes were knocked-in to replace mouse endogenous *disc1* gene. E1 and E2 are exon 1 and 2 of mouse *disc1* gene. **(B)** RT-PCR analysis of the expression of the DB7 fusion genes in humanized DISC1-Boymaw mice (DISC1^H^). The two PCR primers were localized in exon 1 and DB7, respectively (arrows). The expression of the bi-cistronic gene generated a 150 bp cDNA fragment after RT-PCR. Correct splicing was confirmed after sequencing. **(C)** Western blot analysis of the DB7 fusion protein in cultured primary neurons. Expression of the DISC1-Boymaw (DB7) fusion proteins was detected from the primary neurons isolated from the humanized DISC1-Boymaw mice. The 75 kD protein band is very close to predicted molecular size (about 73 kD). **(D)** Immunocytochemical staining of the DB7 fusion proteins in the primary neurons. Primary neurons were isolated from postnatal day 1 mice, and cultured 4 days *in vitro* before immunostaining. Co-localization between MAP2 and DB7 proteins was observed. Scale bar: 20 µm**. (E)** Decreased MTT reduction in brain homogenate of neonatal heterozygous mice (DISC1^H^) (n=14) in comparison with wildtype control littermates (n=12) after normalization with total amount of proteins (t(24)=2.984, p < 0.01, unpaired, two-tailed student’s t-test). **(F)** Reduced expression of Cyb5r3 in the ER isolated from the brains of neonatal DISC1-Boymaw mice (n=16) and wildtype littermates (n=18) after normalization with ER marker Cypor (unpaired, two-tailed student’s t-test, t(32)=2.58, p < 0.05). Error bar: SEM. (* p < 0.05, ** p < 0.01)

### Inhibition of rRNA Expression and Protein Translation *in vivo*

We were particularly interested in the inhibition of both rRNA expression and protein translation in HEK293T cells expressing the DB7 fusion gene. To investigate whether expression of the DB7 fusion gene also decreases rRNA expression *in vivo*, we extracted total RNA from mouse hippocampus and cortex. Heterozygous DISC1-Boymaw mice had significantly less hippocampal rRNA than their wildtype siblings (Figure 6A). No difference in hippocampal DNA was found between the two genotypes. Reduction of rRNA expression was also observed in the cortex of the heterozygous DISC1-Boymaw mice (Figure S6A). To confirm whether rRNA expression is reduced in neuronal cells, we conducted non-radioactive chromogenic RNA *in situ* hybridization to visualize 18S rRNA expression (Figure 6B)(Figure S6B). Reduced expression of 18S rRNA was observed across mouse brain including both hippocampal and cortical neuronal cells in the heterozygous DISC1-Boymaw mice. Consistent with reduction of rRNA expression, we found significant reduction of protein translation in the hippocampus of the heterozygous DISC1-Boymaw mice after intracerebroventricular (ICV) injection of puromycin in *in vivo* SUnSET experiments (Figure 6C, 6D) (24). To investigate whether the inhibition of protein translation is independent of potential differential brain activities, we examined protein translation activities in cultured primary neurons isolated from postnatal day 1 mice. Protein translation activity was significantly lower in the primary neurons isolated from the heterozygous DISC1-Boymaw mice than their wildtype siblings (Figure 6E, 6F).

**Figure 6.**
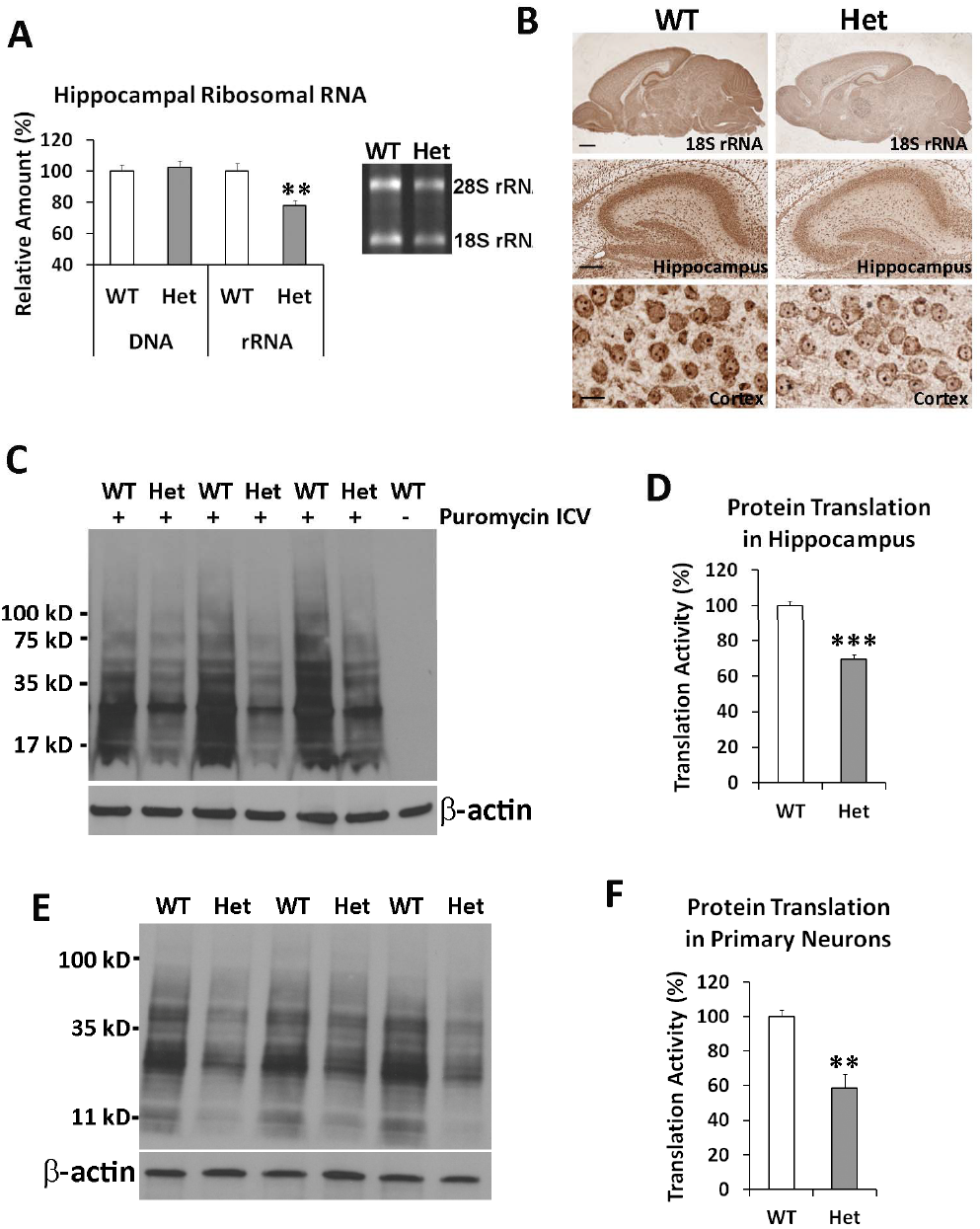
Decreased rRNA Expression and Protein Translation. **(A)** Hippocampal DNA and RNA were isolated from three-week old heterozygous DISC1-Boymaw (n=8) and wildtype control mice (n=12) respectively. No difference in total amount of DNA was observed between the two genotypes. However, significant reduction of total RNA was observed (unpaired, two-tailed student’s t-test, t(18)=3.33, p < 0.01). After gel electrophoresis, 28S and 18S rRNA displayed significant reduction in the hippocampus of the heterozygous DISC1-Boymaw mice. **(B)** Chromogenic RNA *in situ* hybridization of 18S rRNA was conducted in the mouse brains at postnatal day 7 (Scale bar: 400 µm). Reduction of 18S rRNA was observed in both hippocampal neuronal cells (scale bar: 100 µm) as well as cortical neuronal cells (scale bar: 10 µm). Nuclear rRNA was readily detected in the nucleolus of the cortical neurons. **(C)** Protein translation activities in hippocampus. Intracerebroventricular injection of puromycin generated massive labeling of new protein synthesis in mouse hippocampus. Decreased puromycin labeling can be readily observed in the heterozygous mice. **(D)** Significant reduction of protein synthesis activities was found in the hippocampus of the heterozygous DISC1-Boymaw mice in comparison with wildtype sibling mice (unpaired, two-tailed student’s t-test, t(4)=9.28, p < 0.001). **(E)** Primary neurons were isolated from postnatal day 1 mice and cultured at 4 days *in vitro* before *SUnSET* experiments. After pulse-labeling, Western blot revealed less puromycin labeling in the primary neurons from the heterozygous DISC1-Boymaw mice than their wildtype siblings. **(F)** Significant reduction of protein synthesis was detected (unpaired, two-tailed student’s t-test, t(4)=4.74, p < 0.01). Error bar: SEM. (* p < 0.05, ** p < 0.01, *** p < 0.001)

### Decreased Gad67, Nmdar1, and Psd95 Expression

We next examined whether general inhibition of protein translation in the humanized mice reduces protein expression of genes critical for pathophysiology of schizophrenia and other psychiatric disorders. Genes encoding Gad67, Nmdar1, and Psd95 proteins were examined because of their central roles in GABAergic and glutamatergic neurotransmission as well as synaptogenesis. There are also many reports on their down-regulation in psychiatric disorders (25–32), and degradation of PSD95 is proposed to play a key role in autism (33). We first conducted real-time RT-PCR quantification of their RNA expression, and did not find significant reduction in the humanized mice (Figure 7A). To examine their protein expression, we conducted Western blot analyses. Significant reduction of Gad67, Nmdar1, and Psd95 was observed in the heterozygous DISC1-Boymaw mice (Figure 7B). No significant difference was found in synaptophysin. It has been reported that neural circuitry activity can regulate Gad67 gene expression at the system level (34). To rule out any potential effects from differential neural circuitry activity of individual mice, we examined Gad67 expression in primary neuron culture. Reduction of Gad67 proteins was remarkable in the primary neurons isolated from both E18.5 embryos and postnatal day 1 (P1) DISC1-Boymaw mice (Figure 7C). Moderate reduction of Psd95 was also observed. Expression of Nmdar1 was very low and inconclusive. Immunohistochemical analyses were also conducted to examine expression pattern of these proteins in adult mouse brains (Figure 7D). Gad67 and Nmdar1 proteins displayed similar expression patterns between the heterozygous DISC1-Boymaw mice and wildtype siblings. In the heterozygous DISC1-Boymaw mouse brains, there is no obvious loss of Gad67 positive cortical GABAergic interneurons (data not shown). Anti-Psd95 antibody did not work on mouse paraffin sections. We also included anti-parvalbumin antibody in the analysis to examine whether this major subgroup of cortical GABAergic interneurons may be abnormal in the heterozygous DISC1-Boymaw mice. We did not find differences in either the number of positive neurons or intensity of parvalbumin expression between the two genotypes.

**Figure 7.**
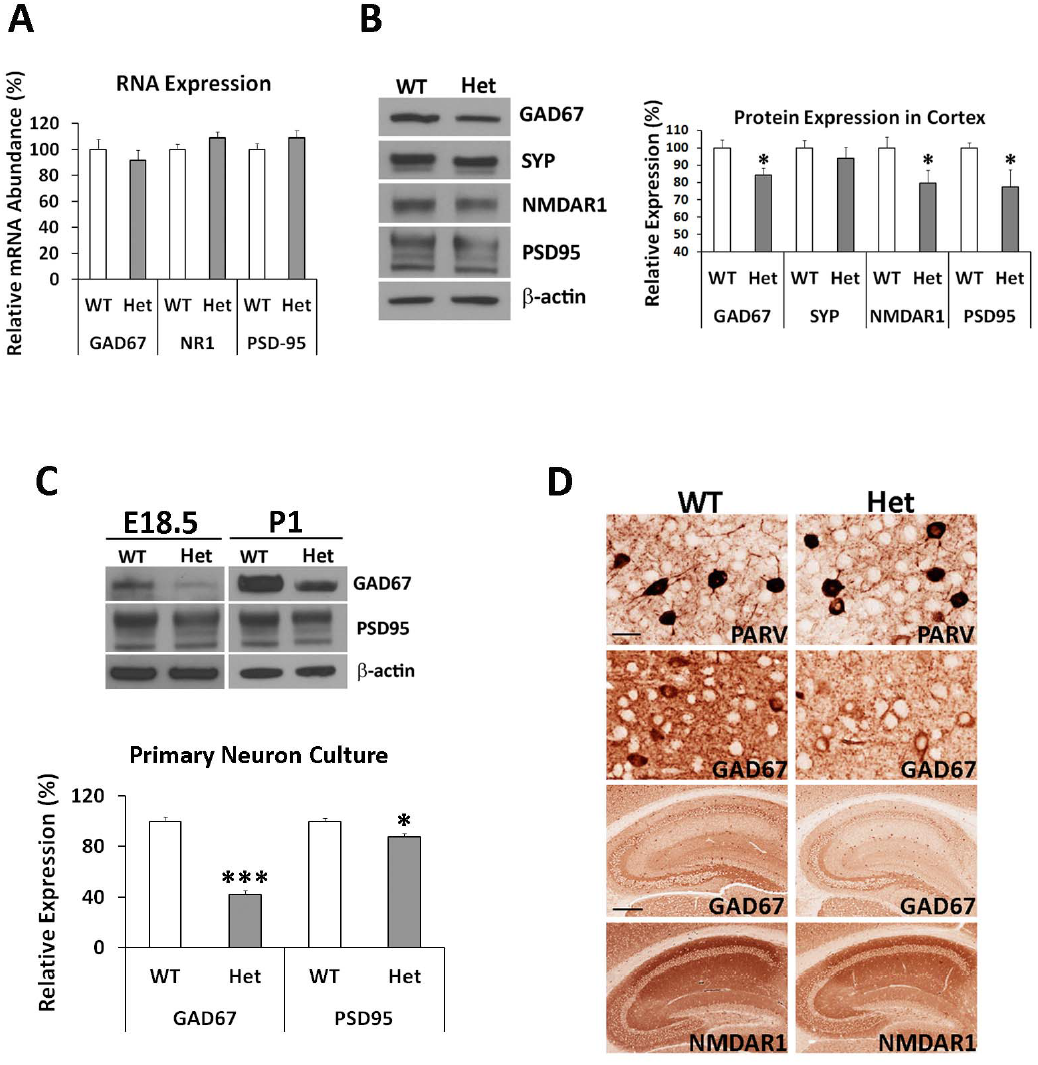
Reduction of Gad67, Nmdar1, and Psd95 Proteins. **(A)** Q-PCR analysis of mRNA transcripts of *Gad67*, *Nmdar1*, and *Psd95* genes in adult mouse brain. There was no significant difference between wildtype and the heterozygous DISC1-Boymaw mice in *Gad67* (unpaired, two-tailed student’s t-test, t(19)=0.73, n.s.), *Nmdar1* (unpaired, two-tailed student’s t-test, t(19)= -1.44, n.s.), and *Psd95* (unpaired, two-tailed student’s t-test, t(19)= -1.17, n.s.) expression. **(B)** Western blot analysis and quantification of Gad67, synaptophysin (Syp), Nmdar1, and Psd95 in the brains of the adult heterozygous DISC1-Boymaw (n=9) and wildtype (n=9) mice. Unpaired two-tailed student’s t-test was used for statistical analysis. No significant reduction of synaptophysin was observed between the two genotypes. Significant reduction of Gad67 (t(16)=2.37, p < 0.05), Nmdar1(t(16)=2.55, p < 0.05), and Psd95(t(16)=2.58, p < 0.05) proteins was observed after normalization with either total amount of loading protein or β-actin expression. **(C)** In primary neurons isolated from cortex and striatum of postnatal day 1 mice, significant reduction of both Gad67 (t(4)=11.92, p < 0.001) and Psd95 (t(4)=3.30, p < 0.05) proteins was observed in cultured primary neurons at 4 days *in vitro* from the DISC1-Boymaw mice (n=3) in comparison with wildtype controls (n=3) (unpaired, two-tailed student’s t-test). Reduction of both proteins was further confirmed in the primary neuron culture (isolated from cortex and striatum) at 4 days *in vitro* from E18.5 embryos. **(D)** Immunohistochemical analysis of parvalbumin, Gad67, and Nmdar1 proteins in the heterozygous DISC1-Boymaw mice (anti-Psd95 antibodies did not work on paraffin sections). Reduction of Gad67 and Nmdar1 was observed in both hippocampus (scale bar: 100 µm) and cortex (scale bar: 20 µm). No difference was detected in parvalbumin expression. White “spots” were nuclei not stained by antibodies. Error bar: SEM. (* p < 0.05, ** p < 0.01, *** p < 0.001).

### Behavioral Analyses of the Humanized DISC1-Boymaw Mice

Since the DB7 fusion gene is a gain-of-function mutation in inhibiting protein translation, the function of the fusion gene was therefore studied in the heterozygous DISC1-Boymaw mice. Mice on the C57 background carry wildtype *Disc1* genes, but 129S mice are natural null mutants of the full length *disc1* gene (6, 35) (Figure S6C, S6D). By following the recommendation of using F1 genetic background by Banbury Conference on Genetic Background in Mice (36), we generated heterozygous DISC1-Boymaw mice on F1 129S/C57 genetic background for behavioral studies (Figure 8A). In the F1 generation, both wildtype and the DISC1-Boymaw heterozygous mice carry one allele of mouse wildtype *Disc1* gene. The DISC1-Boymaw heterozygous mice were healthy and indistinguishable from their wildtype sibling mice. Prepulse inhibition (PPI) and habituation of acoustic startle were examined because they are both deficient in patients with schizophrenia and/or other major psychiatric disorders (37, 38). No differences in PPI (100 msec interstimulus interval; ISI), startle, or startle habituation were found between the genotypes (Figure 8B, 8C). However, we noticed that there was a large startle variation between individuals with a significant number of mice with low startle (below 50)(Figure 8D), which could complicate the measurement of PPI. The source of such a large startle variation remains unknown since there is no genetic segregation in F1 generation. Schizophrenia patients are reported to have high sensitivity and prolonged responses to ketamine, an NMDA receptor antagonist (39–41). Cortical GABAergic inhibitory interneurons have been proposed to be the primary targets of NMDAR antagonists (42, 43). In light of our findings of reduced Nmdar1 and Gad67 proteins, we examined responses of the heterozygous DISC1-Boymaw male mice to ketamine in the mouse Behavioral Pattern Monitor (BPM) (44). A significant gene by ketamine interaction was observed (F(1,27)=4.30, p < 0.05). *Post hoc* analyses revealed that the heterozygous DISC1-Boymaw male mice display a larger increase in total distance traveled (Figure 8E), counts (Figure S7A), and transitions (Figure S7B) than sibling wildtype mice in locomotion after ketamine injection. It is unlikely that less locomotion of the wildtype mice results from ketamine overdose, since we did not observe initial suppression and later increase of locomotion by ketamine overdosing in the wildtype mice. However, multiple dosages of ketamine are needed to fully address the ketamine sensitivity of the heterozygous DISC1-Boymaw mice in the future.

**Figure 8.**
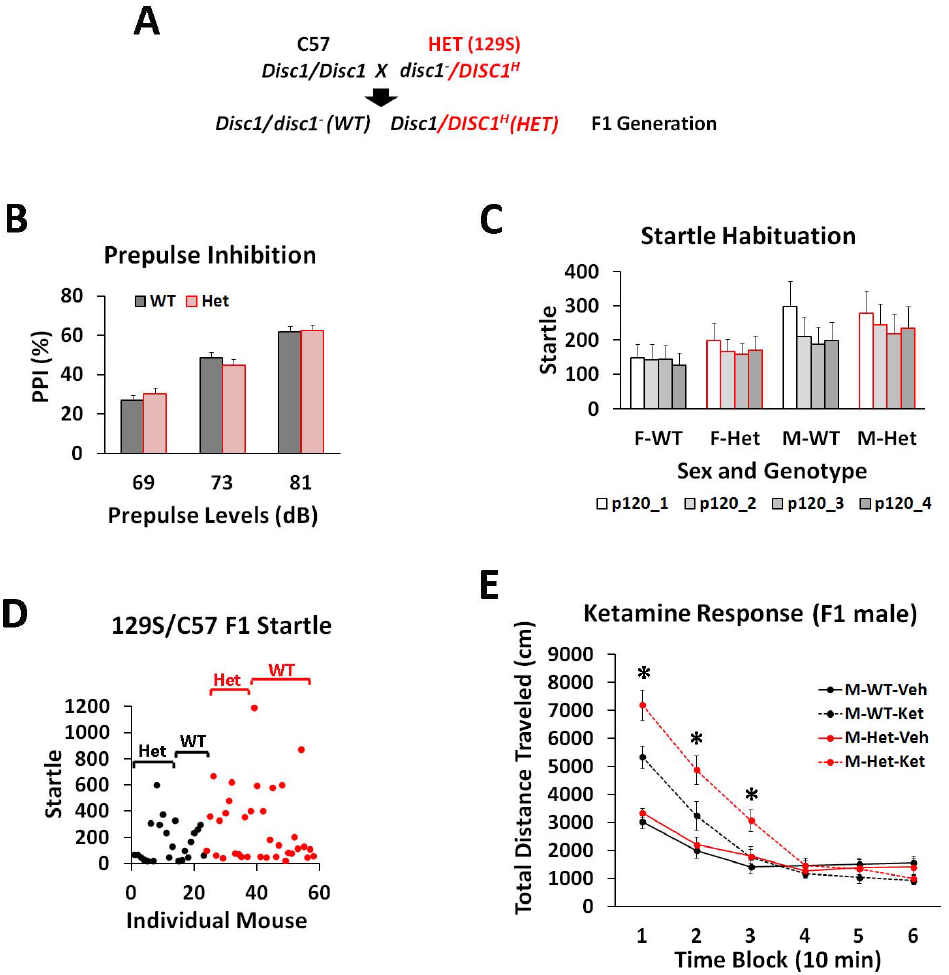
Behavioral Analyses of the Humanized DISC1-Boymaw Mice on F1 129S/C57 Genetic Background. **(A)** The breeding scheme of mice on F1 129S/C57 background. In the F1 generation, there is no genetic segregation between individual mice. Both wildtype and the heterozygous DISC1-Boymaw mice carry one wildtype copy of mouse *Disc1* gene. **(B)** The cohort of F1 mice consists of 28 wildtype and 26 heterozygous DISC1-Boymaw mice. There was no sex effect or gene effect on PPI (F(1,52)=0.00, n.s.). **(C)** There was no gene effect on startle habituation (F(1,50)=0.21, n.s.). **(D)** A large startle variation was observed between individual mice despite the fact that they are genetically identical. Dots represent individual mice (black = females, red = males). **(E)** Males were tested for their responses to ketamine in the BPM. There were 15 wildtype and 14 heterozygous age-matched males. A significant gene by ketaminne interaction was observed (F(1,27)=4.30, p < 0.05). *Post hoc* analyses (Tukey Studentized Range Test) revealed that the heterozygous mice displayed significantly increased responses to ketamine in comparison with wildtype male controls in the first three blocks. Error bar: SEM. # p < 0.10; * p < 0.05; ** p < 0.01.

We next examined the behavioral phenotypes of the humanized mice in the absence of full-length wildtype *Disc1* gene on 129S genetic background (Figure 9A). Because wildtype 129S mice carry mutant *disc1* genes, we define that “wildtype mice” in our studies refer to mice that do not carry the knocked-in DISC1-Boymaw fusion gene, rather than refer to the status of mouse endogenous *disc1* gene. Both wildtype and heterozygous mice were healthy; however, male heterozygous mice were slightly, but significantly, smaller than their wildtype littermates (Figure 9B). No significant body weight difference was observed in females between the two genotypes. In contrast to mice on mixed 129S/C57 genetic background, mice on 129S pure genetic background have higher startle with less variation between individuals (Figure S8A), which makes the 129S genetic background more suitable for PPI studies. To extend the PPI studies with ISI at 100 msec in the F1 mice, we performed PPI tests with various ISIs in this cohort. When PPI was conducted with a short ISI at 25 msec, however, enhanced PPI, or impaired prepulse facilitation, was observed in the heterozygous DISC1-Boymaw mice, suggesting abnormal information processing (Figure 9C). After ketamine administration in the BPM, significant gene (F(1,52)=11.42, p < 0.01), ketamine (F(152)=24.06, p < 0.0001), and gene X ketamine X sex interaction (F(1,52)=5.40, p < 0.05) effects were observed in total distance traveled. In male mice, a significant gene X ketamine interaction (F(1,26)=8.61, p < 0.01) was observed in the distance traveled (Figure 9D) and other measures of locomotion activity (Figure S8B, S8C, S8D). *Post hoc* analyses revealed significantly increased locomotion in the heterozygous DISC1-Boymaw mice injected with ketamine in all blocks except the first and last, suggesting increased and prolonged responses to ketamine. In female mice, a significant gene effect (F(1,26)=5.63, p < 0.05) was detected without gene X ketamine interaction (Figure S8E). To examine whether the heterozygous mice display behavioral phenotypes related to depression, a tail suspension test was conducted to assess mouse behavioral despair (45, 46). The heterozygous

**Figure 9.**
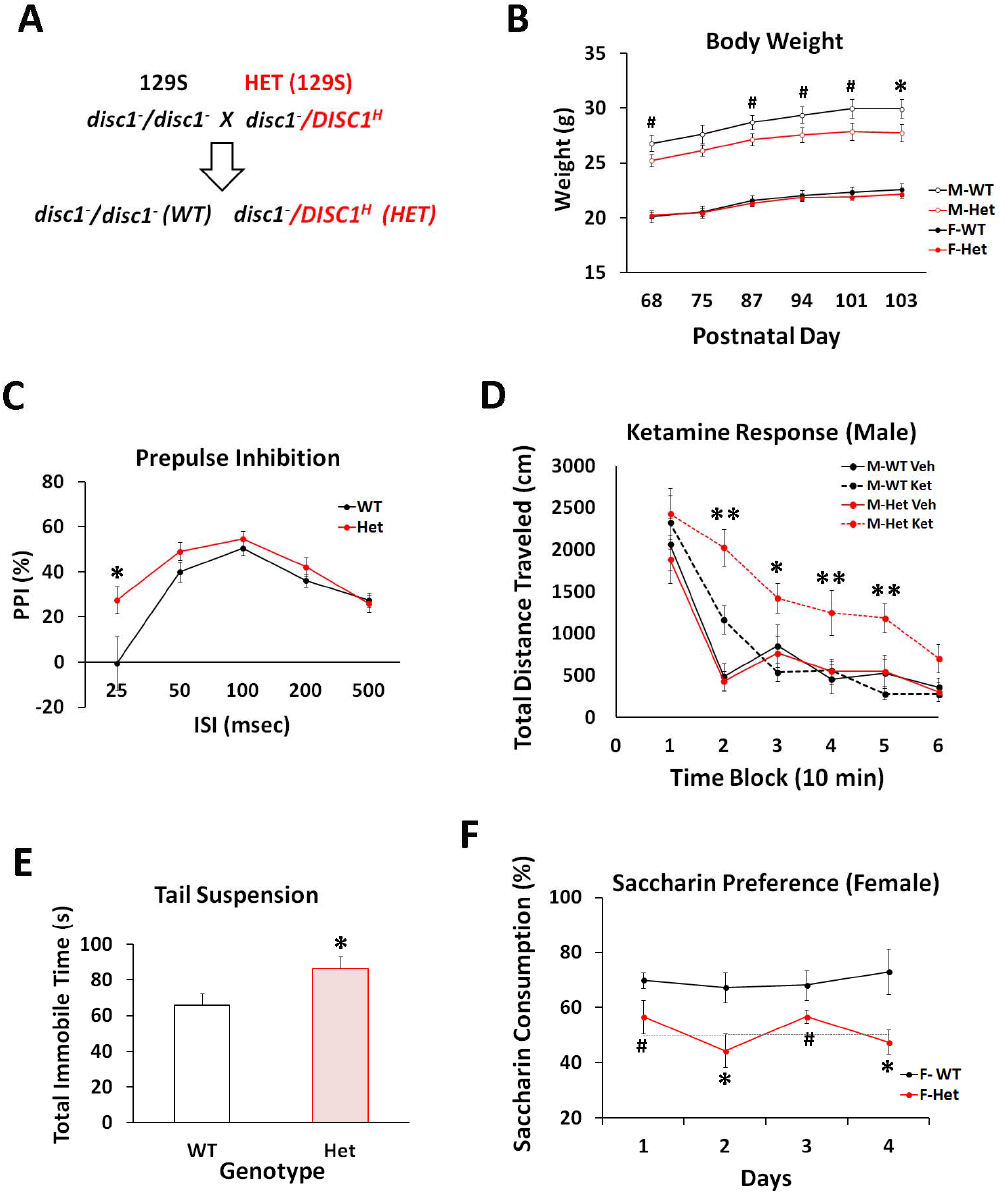
Behavioral Characterization of the Humanized DISC1-Boymaw Mice on 129S Background. **(A)** Breeding of wildtype and the heterozygous DISC1-Boymaw mice on 129S pure genetic background. 129S mice carry a frame-shift mutation in exon 6 of the *disc1* gene. **(B)** The cohort consisted of 14 wildtype females, 14 heterozygous females, 13 wildtype males, and 15 heterozygous males. The age range of the birth date between individual mice was less than a week. Body weight was compared between the wildtype and the heterozygous mice. Heterozygous males displayed slightly, but significantly, reduced body weight at postnatal day 103 (unpaired, two-tailed student’s t-test, t(26)=2.132, p < 0.05). No difference was found in body weight between female wildtype and heterozygous mice. **(C)** There was a significant gene effect (F(1,54)=4.41, p < 0.05) in PPI at 69 dB prepulse. A significant gene X ISI interaction was also observed (F(4,216)=2.90, p < 0.05). *Post hoc* analysis revealed that the heterozygous DISC1-Boymaw mice displayed increased PPI (or impaired prepulse facilitation) with ISI at 25 msec. No sex difference was observed. **(D)** The male heterozygous DISC1-Boymaw mice display a significant gene effect (F(1,26)=6.26, p < 0.05) as well as a gene X ketamine interaction (F(1,26)=8.61, p < 0.01). *Post hoc* analyses (Tukey Studentized Range Test) revealed that the male heterozygous mice displayed significantly increased and prolonged responses to ketamine in comparison with wildtype male controls in all time blocks except the first and last one. **(E)** Total immobility time (seconds) was recorded during a 6 min suspension test. Five wildtype male mice were lost because of fighting in cages after previous tests. The heterozygous DISC1-Boymaw mice (n=29) displayed significantly more immobility time than the wildtype control mice (n=22) (unpaired, two-tailed student’s t-test, t(49)=2.22, p < 0.05). **(F)** 2 or 3 mice were grouped in a single cage according to genotype. Saccharin preference test was conducted for 4 consecutive days. Consumption of saccharin was measured each day, and compared between the wildtype and the heterozygous mice in each sex. Significant gene effect was detected (F(1,11)=17.85, p < 0.01) between female wildtype (n=7) and the DISC1-Boymaw heterozygous mouse cages (n=6). *Post hoc* analysis revealed that the female heterozygous mice displayed significantly less saccharin consumption than female wildtype controls mice at day 2 and day 4. Error bar: SEM. # p < 0.10; * p < 0.05; ** p < 0.01.

DISC1-Boymaw mice displayed significantly more immobility time than wildtype sibling controls without sex effects (p < 0.05)(Figure 9E). A saccharin preference test was also conducted to examine anhedonia-like behavior, as seen in depression in humans, in the heterozygous mice. Wildtype female mice displayed significantly higher preference for saccharin than the heterozygous female mice (Figure 9F). Because wildtype male mice had no significant preference for saccharin, the heterozygous DISC1-Boymaw male mice cannot be assessed for depression-like behavior using this behavioral paradigm (Figure S8F). It has been reported that some rodent strains display a sex difference in saccharin preference in which females have strong preference while males have little preference or even some avoidance (47). In general, 129S mice are reported to have only slight saccharin preference (http://www.criver.com/SiteCollectionDocuments/129.pdf). These findings may explain why wildtype male mice on the 129S background did not display saccharin preference. Taken together, the humanized DISC1-Boymaw male mice consistently displayed a prolonged response to ketamine on different genetic backgrounds, mimicking phenotypes of schizophrenia and potentially depression. Replication of this phenotype in different cohorts of mice helped rule out type I error by multiple comparisons. Abnormal information processing of startle and depressive-related behaviors have also been suggested, and further confirmation is needed in future.

### Abnormal Erythropoiesis in Humanized DISC1-Boymaw Mice

Mutations of ribosomal protein genes, which impair protein translation, have been demonstrated to cause Diamond-Blackfan anemia (48). Since the human DISC1 gene is also expressed in lymphoblastoid cells, we conducted hematological analysis of the heterozygous DISC1-Boymaw mice to investigate whether inhibition of protein translation by the DISC1-Boymaw fusion gene may cause anemia (Figure 10A). Interestingly, 2 out of 10 heterozygous DISC1-Boymaw male mice displayed lower concentrations of hemoglobin (Figure 10B)(Table S2), indicating different degrees of anemia (49). A significant difference between the two genotypes is an increase of red blood cell distribution width (RDW), which measures the variation of red blood cell volume, in the heterozygous DISC1-Boymaw mice (Figure 10C). We re-examined RDW in some of these mice one week later to investigate RDW stability. RDW displayed little change in wildtype mice, but higher RDW fluctuation was observed in their sibling heterozygous DISC1-Boymaw mice (Figure 10D). These data indicated that the DISC1-Boymaw fusion gene impairs the stability of erythropoiesis in the heterozygous DISC1-Boymaw mice.

**Figure 10.**
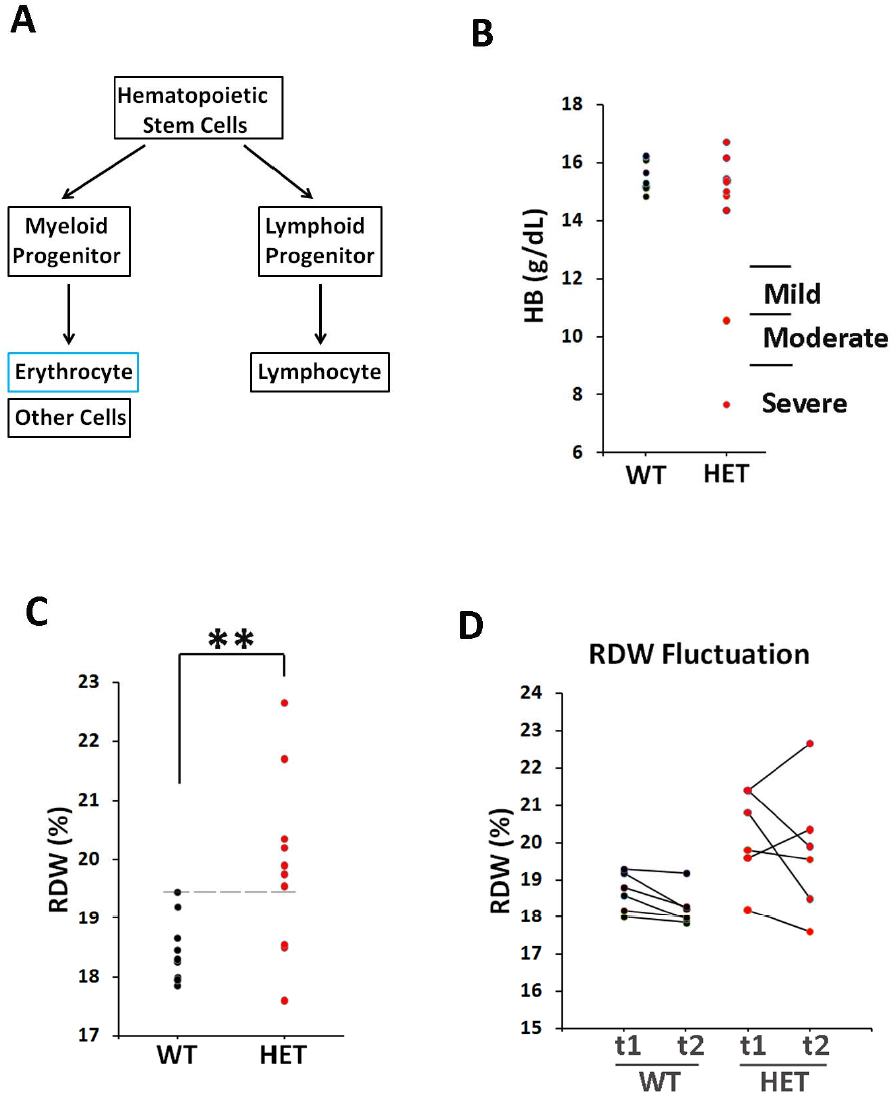
Increased RDW in the Humanized DISC1-Boymaw Mice. **(A)** Schematic pathway of erythropoiesis. **(B)** Concentration of hemoglobin (HB) was measured in the blood of individual male mice. Severe, moderate, and mild anemia was diagnosed according to hemoglobin concentrations (49). Two heterozygous DISC1-Boymaw mice suffered from severe and moderate anemia. All wildtype mice were normal. Each dot represents an individual mouse. **(C)** Homogeneity of variances of RDW data was first assessed with Levene’s test (p = 0.057). Significantly higher RDW was found in the heterozygous DISC1-Boymaw mice (F(1,19)=8.451, ** p < 0.01). Most of these mice have RDW above the upper limit of the normal RDW range of their wildtype sibling mice (dashed line). **(D)** 6 wildtype and 6 heterozygous DISC1-Boymaw male mice were re-examined one week later (t2) for their RDW stability, and compared with their initial (t1) RDW measurement. There was little change in RDW of the wildtype mice during this period. However, high magnitude of RDW fluctuation was observed in the heterozygous DISC1-Boymaw mice, indicating unstable erythropoiesis.

## Discussion

The t(1;11) chromosome translocation in the Scottish family co-segregates with schizophrenia and major depression with 70% penetrance (1–3). Despite extensive studies on the DISC1 gene, the role of the disruption of the DISC1 gene alone remains unclear in the pathogenesis of major psychiatric disorders (35, 50, 51). Our previous studies revealed that the DISC1 gene disruption is accompanied by disruption of the Boymaw gene and generation of the DISC1-Boymaw and Boymaw-DISC1 fusion genes (10, 11). In the present study, we found that expression of the DB7 fusion gene decreases NADH oxidoreductase activities, rRNA synthesis, and protein translation in both *in vitro* cell studies and *in vivo* humanized DISC1-Boymaw mice. Despite inhibition of protein translation by the DB7 fusion gene, however, we did not find gross anatomical abnormalities in the brain of the heterozygous DISC1-Boymaw mice.

Inhibition of protein translation does not always cause obvious developmental defects. For example, reduced protein translation by silencing translational initiation factor eIF4E isoform (IFE-2) has no effects on *C. elegans* development and body size at 20 °C (52). Even though we did not find obvious developmental abnormalities, reduced expression of Gad67, Nmdar1, and Psd95 proteins was observed in the brain of the heterozygous DISC1-Boymaw mice. Behavioral studies showed that the humanized mice display phenotypes related to both schizophrenia and depression. Our studies therefore suggest that the DB7 fusion gene could be one of the causal gene mutation(s) in the pathogenesis of major psychiatric disorders in the Scottish family. It should be kept in mind, however, that the humanized mice also harbor the BD13 fusion gene in addition to the DB7 fusion gene. Reduction of protein translation in the brains of the humanized mice is likely caused by the expression of the DB7 fusion gene, according to our *in vitro* cell transfection studies. However, we cannot rule out a potential contribution from the BD13 fusion gene to the behavioral phenotypes, although we did not find any cellular phenotypes in HEK293T cells expressing the BD13 fusion gene.

Our *in vitro* studies revealed that the DB7 fusion protein is unstable and its expression decreases intracellular oxidoreductase activities, rRNA synthesis, and protein translation. These phenotypes are unlikely caused by the insolubility of the DB7 proteins *per se*, since the much more abundant insoluble DISC1 and truncated DISC1 proteins did not generate any of these cellular phenotypes in the over-expressing HEK293T cells. However, the insoluble DISC1 proteins in HEK293T cells may be structurally different from the insoluble proteins found in postmortem human brains (12). Surprisingly, fusion of the Boymaw gene to a randomly selected gene (fluorescence marker) generated the same cellular phenotypes as the DB7 fusion gene. It will be interesting to know whether the human Boymaw gene encodes any protein or peptide. Regardless of the Boymaw gene being a coding or non-coding RNA, the DB7 fusion gene appears to be a gain-of-function mutation. The role of the Boymaw gene merits further studies. Unfortunately, there is no human Boymaw gene orthologue in mouse genome.

Eykelenboom et al identified three different fusion transcripts from the patient lymphoblastoid cells from the Scottish family (53). The DB7 fusion gene used in our studies encodes exactly the same fusion protein as their CP60 transcript. The second CP69 fusion transcript is generated by alternative splicing 12 nucleotides upstream of the stop codon of the CP60 transcript. Our deletion analysis demonstrated that the last few amino acid residues of Boymaw are not required for inhibition of MTT reduction. Therefore, it is likely that the CP69 fusion protein is also capable of inhibiting MTT reduction. The third CP1 fusion transcript encodes a protein with just one more amino acid addition to the C-terminus of the truncated DISC1 protein that has been used in the generation of several mouse DISC1 models (54, 55). The CP1 protein does not display any detectable toxicity to mitochondria, which is consistent with our observation that over-expression of truncated DISC1 proteins has no effect on MTT reduction, rRNA expression, and protein translation. Our immunocytochemical analysis confirmed localization of the DB7 fusion protein (but not DISC1-FL, truncated DISC1, or BD13) in mitochondria (Figure S9). Interestingly, MTT reduction was also reduced in mitochondria of cells expressing the DB7 fusion gene (Figure 3C). Decreased activities of mitochondrial NADH oxidoreductases may relate to loss of mitochondrial membrane potential in cells expressing the DB7 fusion protein (53). We therefore hypothesize that reduction of oxidoreductase activity may decrease cell metabolism to down-regulate rRNA expression and protein translation. One potential pathway could be that inhibition of the mitochondrial oxidoreductase activities reduces ATP production (Figure 11), which is supported by the finding of loss of mitochondrial membrane potential in cells expressing the DB7 fusion protein (53). Reduction of ATP can trigger activation of the AMP-activated protein kinase (AMPK), a sensor of cell energy status (56). The AMPK pathway is a well established pathway to regulate protein translation. In this pathway, the activated AMPK phosphorylates TSC1/TSC2 to inhibit mTOR. mTOR kinase plays a central role to stimulate protein translation either directly via activation of translational initiation factors (57) or indirectly via phosphorylating transcription initiation factor TIF-1A to promote rRNA transcription (58). Mutation of TSC1/TSC2 leads to autism (59), which shares many common psychopathological features with major psychiatric disorders. Another potential pathway linking cellular metabolism to rRNA transcription could be initiated through the Sirtuin family of proteins, which are NAD^+^-dependent deacetylase to sense cellular NAD^+^ changes. SIRT1 (Sirtuin 1) has been demonstrated to epigenetically silence rDNA chromatin (60, 61). However, it is also reported that nucleolar SIRT7 (Sirtuin 7) functions as an activator for rRNA transcription (62). It is clear that the potential underlying mechanisms are complex and much more work is needed.

**Figure 11.**
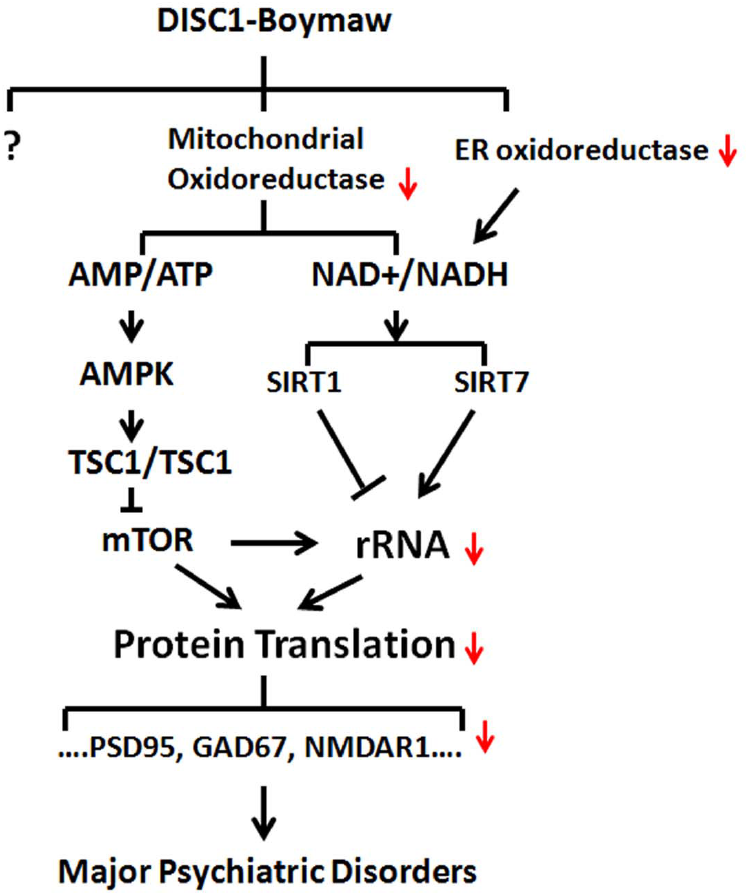
Hypothetical Pathways of the DB7 Fusion Gene. Cell metabolism is tightly coupled with protein synthesis. Reduction of NADH oxidoreductase activity may alter production of ATP and NAD^+^ which can trigger at least two known pathways (AMPK and Sirtuin) to down-regulate rRNA transcription and protein translation. AMPK: AMP-activated protein kinase. TSC1 and TSC2: tuberous sclerosis complex 1 and 2. mTOR: mammalian target of rapamycin.

One of the key questions is how expression of the DB7 fusion protein reduces oxidoreductase activity. Identification of these NADH oxidoreductases in mitochondrion and cytoplasm is critical to unravel the signaling pathways from cellular metabolic activity to protein translation. In contrast to inhibition of protein translation by the DB7 fusion gene, exaggerated protein translation has been suggested as the core pathophysiology of Fragile X Syndrome (63) and other autism genes (24). It is possible that an optimal protein translation is necessary for neural plasticity (64), and excessive or less protein translation are harmful to brain functions. Translational control of protein synthesis may play important roles in many other psychiatric disorders. Indeed, it was reported that hypermethylation of ribosomal RNA gene promoter was associated with suicide (65), mild cognitive impairment and Alzheimer’s disease (66), as well as Huntington’s diseases (67). Considering that rRNA synthesis is a rate-limiting step of ribosome biogenesis (68), reduction of rRNA will likely result in a decrease of protein translation. Our SUnSET experiments clearly demonstrated inhibition of protein translation in both the cells transiently expressing the DB7 fusion gene and the heterozygous DISC1-Boymaw mice. Since we did not find significant reduction of mRNA transcripts of *Gad67*, *Nmdar1*, and *Psd95* genes, the reduction of general protein translation therefore could be responsible for the decrease of Gad67, Nmdar1, and Psd95 protein expression in the heterozygous DISC1-Boymaw mice. Indeed, it was reported that cellular abundance of proteins is predominantly controlled at the level of protein translation (69, 70). However, it is also possible that protein turnover may be altered and contributes to the reduction of these proteins in the humanized mice. The effects of the expression of the DB7 gene may not be restricted to the inhibition of protein translation. For example, we observed reduced expression of Cyb5r3 reductase in endoplasmic reticulum (ER), where the enzyme plays an important role in lipid biosynthesis and has been demonstrated to be essential for brain development (71–73).

Impaired NMDAR-mediated neurotransmission has been well established in the pathophysiology of schizophrenia. Prolonged ketamine effects and exacerbation of symptoms are observed in schizophrenia patients (39–41) and our mouse genetic model (74). Consistent with the findings in human patients, we found reduction of both Nmdar1 and Gad67 proteins in heterozygous DISC1-Boymaw mice which display prolonged responses to the noncompetitive NMDAR antagonist, ketamine. This phenotype has been replicated in different cohorts of mice on various genetic backgrounds regardless of the presence or absence of wildtype *Disc1* gene. It is unknown why the female humanized mice do not show prolonged responses to ketamine. The female humanized mice are not symptom-free, since they display abnormal phenotypes in other behavioral paradigms. We found increased PPI in the heterozygous DISC1-Boymaw mice at a short ISI (25 msec) in both sexes. In mice, PPI diminishes with decreasing ISI starting from 100 msec, and eventually transforms into prepulse facilitation. Abnormally enhanced PPI at short ISI is indicative of altered information processing in the heterozygous DISC1-Boymaw mice. Although this alteration in PPI differs from that seen in patients with schizophrenia (75), enhanced PPI at short ISI has been reported in rats treated with dopamine receptor agonists (76, 77).

Inhibition of protein translation by the fusion gene impacts not only the brain but also erythropoiesis of the heterozygous DISC1-Boymaw mice. We found an incomplete penetrance of the DISC1-Boymaw fusion gene to anemia, consistent with the role of impaired protein translation demonstrated by ribosomal protein gene mutations in Diamond-Blackfan anemia (48). Interestingly, the heterozygous DISC1-Boymaw mice also display increased RDW. In human, over 2 million red blood cells are generated every second with small variation in cell volume (human RDW: 11.5–14.5%). It is possible that such an extremely active and precise biological process in mouse is highly susceptible to the DISC1-Boymaw mutation which confers even mild inhibition of protein translation. Recently, an association between increased RDW and depression was reported from a multi-year large scale study in humans (78). The study found a gradient positive correlation between increased RDW and the risk of developing depression in thousands of patients. It will be interesting to know whether patients with major depression in the Scottish family may display increased RDW.

It should be kept in mind that our studies are focused on the investigation of the biological functions of the DISC1-Boymaw fusion genes. To that end, the 129S genetic background, which contains a frame-shift mutation in endogenous *disc1* gene, provides a unique advantage for our studies to avoid *Disc1* gene dosage difference between wildtype and the heterozygous DISC1-Boymaw mice. However, the chromosome translocation in the Scottish family generates not only the fusion genes but also loss of one allele of the *DISC1* gene. Therefore, our humanized DISC1-Boymaw mice on 129S background should be viewed as a genetic model for the DISC1-Boymaw fusion genes rather than the chromosome translocation in the Scottish family. In the future, we could generate mice carrying full-length human *DISC1* genes on mouse *Disc1* locus using the *Cre* recombinase. After breeding these *DISC1* mice with the heterozygous DISC1-Boymaw mice, we could generate wildtype mice carrying two human *DISC1* (*DISC1/DISC1*) genes and heterozygous mice carrying one human *DISC1* and one DISC1-Boymaw fusion gene (*DISC1/DISC1^H^*). Comparisons between wildtype (*DISC1/DISC1*) and the heterozygous (*DISC1/DISC1^H^*) will provide a more complete model for the chromosome translocation in the Scottish family.

### Materials and Methods

#### DNA constructs

Full length DISC1, truncated DISC1, DB7, and BD13 fusion genes were generated through PCR amplification and ligation. All expression constructs were tagged with HA epitopes as described (11). The AAV shuttle and pTimer plasmid vectors were used to generate expression constructs, bi-cistronic constructs connected with the IRES of EMCV virus (79), and fluorescence timer (FT) fusion protein constructs, respectively. Boymaw was also fused with FT in pTimer plasmid vector. All constructs were confirmed with complete sequencing. A series of FT-Boymaw C-terminal deletion were generated by using PCR amplification and subsequent In-Fusion ligation (Clontech). The following pairs of PCR primers were used: C1-del: Boymaw-C1-del-f: 5’CCTTGAGTAC CCATACGACG TTCCAGACT3’; Boymaw-C1-del-r: 5’TATGGGTACT CAAGGCTTCT ATCTGTGGT3’; C2-del: Boymaw-C2-del-f: 5’ TCTACCATAC CCATACGACG TTCCAGACT3’; Boymaw-C2-del-r: 5’ TATGGGTATG GTAGAAGCCT GCAGACAAAG3’; Boymaw-C3-del-f: 5’ GAAGTCATAC CCATACGACG TTCCAGACT3’; Boymaw-C3-del-r: 5’ TATGGGTATG ACTTCAGGTA ACTGAAACC3’; Boymaw-C4-del-f: 5’ GAGACATTAC CCATACGACG TTCCAGACT3’; Boymaw-C4-del-r: 5’ TATGGGTAAT GTCTCTTATT CCAGTCTGT 3’.

#### In vitro cell culture

HEK293T, COS-7, and SH-SY5Y cells were cultured in DMEM or DMEM/F12 medium supplemented with 10% fetal bovine serum, penicillin, and streptomycin (Life Technologies, CA) at 37°C in a humidified atmosphere containing 5% CO_2_. HEK293T, COS-7 cell transfections were performed using Mirus TransIT^®^ reagent (Mirus, WI), according to manufacturer’s instructions. SH-SY5Y cells were transduced with recombinant AAV virus. Primary culture of mouse neuronal cells was established from both wildtype and heterozygous DISC1-Boymaw embryos at E18.5 and P1 as described in our previous studies (80). The mouse cerebrum were dissected and cut into small pieces in Neurobasal A medium on ice. The tissue was triturated ten times with a fire polished 9-inch Pasteur pipette and allowed to settle on ice for 1 min. The supernatant was then transferred to a new tube and centrifuged gently at 600–700rpm for 5 min to pellet the cells. The cells were resuspended in B27/Neurobasal medium (B27/Neurobasal with 0.5 mM glutamine, no glutamate, 5 ng/ml FGF2).

#### MTT reduction assays

HEK293T cells and COS-7 cells were seeded at cell density of 1 × 10^5^ cells per well in 24-well plates, 24 hr prior to transfection. Cell proliferation was measured by counting cell numbers with 0.04% Trypan blue 48 hr after transfection. MTT (3-(4, 5-dimethylthiazol-2-yl)-2, 5-diphenyl tetrazolium bromide reduction assay was modified from Mosmann T (81). MTT (Sigma, MO) was dissolved in 1xPBS at the concentration of 5 mg/ml, and filtered through 0.2 um membrane. The stock solution was stored at -20°C. For MTT reduction in living cells, MTT stock solution was added at the final concentration of 0.5 mg/ml, and the cells were incubated at 37°C for 1.25 h. For MTT assay in cell lysate, transfected cells were first lysed in lysis buffer (18) (100 mM NaCl, 10 mM Tris, pH 7.5, 1 mM EDTA, 0.01%Triton X-100, protease inhibitors cocktail) at 4 °C for 30 min with gentle shaking. MTT was added to the lysate at the final concentration of 0.5 mg/ml, and the reaction was incubated at 37 °C for 1 hr. For MTT assay in purified ER and mitochondria, NADH (Sigma, MO) was added in the assay at the final concentration of 0.5 mM, and the reactions were incubated at 37 °C for 10 min for mitochondria, and 30 min for ER, respectively. For the reversal of deficient MTT reduction, 2.5 Unit of E coli ADH (Cat# 49641, Sigma, MO) was added into the DISC1-FL and DB7 cell lysate, respectively, and further incubated at 37 °C for 2 hr. MTT crystal were collected and dissolved in DMSO. Absorbance at 540 nm was measured with SpectraMax M5 Multi-Mode Microplate Readers (Molecular Devices, PA). Neonatal mouse brains were dissected on postnatal day 1. After washing three times with pre-chilled PBS, brains were cut into small pieces in 1.5 ml isotonic extraction buffer (IEB) (10 mM HEPES, pH 7.8, with 0.25 M sucrose, 1 mM EGTA, and 25 mM KCl) containing protease inhibitor cocktail (Sigma, MO), and further homogenized with Dounce homogenizer on ice. Equal amounts of protein homogenate were used for MTT reduction assays.

#### Western blot analysis

Supernatant soluble proteins were extracted from HEK293T cells and COS-7 cells in Passive Lysis Buffer (PLB) (Promega, WI) containing 0.2% Sarkosyl and 1X protease inhibitor cocktail (Sigma, MO) at room temperature for 15 min on a shaker. Total proteins from cell pellets, endoplasmic reticulum, mitochondria, and brain homogenate of neonatal mice were solubilized with sonication in PLB containing 1X protease inhibitor cocktail. Protein concentration was measured with Bradford (Abs 595nm) method using Coomassie Plus Protein Assay (Thermo Scientific, IL). After electrophoresis, proteins were transferred onto PVDF membranes which were subsequently blocked with 5% nonfat dry milk in TBST buffer (pH 7.5, 10 mM Tris-HCl, 150 mM NaCl, and 0.1% Tween 20) at room temperature for 1 hr. The membranes were incubated with the following primary antibodies at 4 °C overnight: mouse monoclonal anti-cytochrome C (1:5000), ab110325 (Abcam, MA); rabbit polyclonal anti-Cyb5r3 (1:600), sc-67284; mouse monoclonal anti-Cypor (1:400); sc-25270; mouse monoclonal anti-β-actin (1:5,000); sc-47778 (Santa Cruz Biotechnology, CA). After washing three times, the membranes were further incubated with horseradish peroxidase (HRP)-conjugated anti-mouse IgG (1:5,000, Cell signaling, MA), anti-rabbit IgG (1:5,000, Santa cruz, CA) for 1.5 hr at room temperature. Rat monoclonal anti-HA antibody (1:500) conjugated with peroxidase (3F10) (Roche, CA) was used to directly detect HA epitope. Quantification of protein expression was conducted with Image J.

#### Proteomic analysis

HEK293T cells were transfected with DISC1-FL and DB7 genes, respectively. Two days later, cells were harvested and sent to UCSD Mass Spectrometry Facility Core for iTRAQ analysis. The samples were labeled with 114 Da, 115 Da, 116 Da, and 117 Da mass tags respectively. Protein identification and quantitation were performed using the Paragon™ Algorithm in ProteinPilot™ Software, searching against the NCBInr database.

#### NADH concentration

Intracellular NADH extraction was conducted as described in Cyclex NAD+/NADH colorimetric Assay Kit (CycLex, Japan). In brief, 2×10^5^ cells were seeded in each well of a12-well plate one day before transfection. Cells were collected 48 hr post-transfection. NADH extraction buffer (50 mM NaOH and 1mM EDTA) was added to cell pellets and gently vortexed 2–3 times. To reduce viscosity, samples were incubated at 60 °C for 30 min. Equal volume of neutralization buffer (0.3 M potassium phosphate buffer, pH7.4) was added and incubated at least 5 min on ice. NADH was extracted from the supernatant solution after spinning down the neutralized samples at 15,000 rpm for 5 min at 4 °C. The cell pellets were solubilized to measure protein concentration for the normalization of NADH concentration. E coli alcohol dehydrogenase (Sigma) was used to measure NADH concentration in MTT reduction assays.

#### Density-gradient ultracentrifugation

Mitochondria and ER were purified from HEK293T cells and neonatal mouse brains via density-gradient ultracentrifugation as described in Endoplasmic Reticulum Isolation Kit (Sigma, MO). In brief, HEK293T cells (about 5 × 10^7^) were incubated in hypotonic buffer (10mM HEPES pH7.8, 1 mM EGTA and 25 mM KCl) for 20 min at 4 °C. The cells were pelleted with 600x g for 5 min, and resuspended with appropriate volume of the isotonic extraction buffer (IEB) (10 mM HEPES, pH 7.8, with 0.25 M sucrose, 1 mM EGTA, and 25 mM KCl). For mouse brains, dissected brains were cut into small pieces in the isotonic extraction buffer (IEB). The cells and the tissue were further homogenized with Dounce homogenizer on ice with 2 ml of IEB. Large cellular debris and nuclei were pelleted after centrifuging at 1,000 x g for 10 min at 4 °C. The supernatant was transferred into a new centrifuge tube and further centrifuged at 12,000 x g for 15 min at 4 °C. Crude mitochondria were pelleted, and the supernatant fraction containing ER was transferred to a new ultracentrifuge tube and centrifuged at 150,000 x g in a Beckman Optima™ LE-80K (Beckman Coulter, CA) at 4 °C for 1 hr to purify ER. ER pellets were resuspended in IEB containing protease inhibitors. Crude mitochondrion pellet was further purified as described (82). In brief, mitochondrion pellet was resuspended in 1 ml of mitochondrion isolation buffer (MIB) (70 mM sucrose, 210 mM D-mannitol, 1 mM EDTA-K_2_, 0.23 mM PMSF, 5 mM Hepes-Tris, pH 7.4, and protease inhibitors cocktail) and layered onto 1.5 ml of 7.5% (w/v) Ficoll-sucrose medium (7.5% Ficoll-MIB) on top of 1 ml of 10% (w/v) Ficoll-sucrose medium (10% Ficoll-MIB) and centrifuged in a Beckman Optima™ LE-80K (Beckman Coulter, CA) for 1.5 hr at 100,000 x g at 4 °C. Mitochondria were pelleted at the bottom of the ultracentrifuge tube. The pellets were suspended in 1 ml of MIB and then centrifuged for 5 min at 15,000 x g at 4 °C. The mitochondrion pellets were washed once in MIB and finally resuspended in appropriate volume of MIB. The purified mitochondria were either used for MTT reduction or stored at -80 °C for subsequent analyses.

#### SUnSET

HEK293 cells seeded at cell density of 1 × 10^5^ cells per well in 24-well plates, 24 hr prior to transfection. After transfection, cells were pulse-labeled with puromycin at the concentration of 10 ug/ml for 10 min in the presence or absence of cycloheximide at final concentration of 25 uM. Total proteins were extracted and load on PAGE gel (10 ug per lane) for Western blot analysis. Anti-puromycin antibody (1:10,000)(12D10, Millipore) was used to detect incorporation of puromycin in newly synthesized proteins. Blots were stripped later and re-probed with anti-β-actin antibody. Image J was used to quantify the incorporation of puromycin.

#### Generation of humanized DISC1-Boymaw mice

Both DB7 and BD13 fusion genes were connected with IRES of EMCV virus (11, 79). The bi-cistronic gene was terminated with SV40 poly (A). A splicing acceptor was added to generate in-frame splicing of the bi-cistron with the first exon of human DISC1 gene. The bicistron gene cassette was flanked with two *loxP* sites. Downstream of the bi-cistron, human DISC1 cDNA was capped with a splicing acceptor. Such a design aimed for conditional restoration of DISC1 gene expression in genetic rescue experiments in the future. Two homology arms with respective sizes of 4 kb and 3 kb were amplified from the mouse disc1 gene of 129S ES cells (129S2/SvPasCrl) to assemble the final gene targeting construct. The final gene targeting construct was electroporated into mouse embryonic stem (ES) cells. After positive and negative selections with G418 and ganciclovir, about 1500 ES cell colonies were picked for further screening. Two homologous recombinant ES cell colonies were identified. After blastocyst injection, chimeric mice were obtained and bred with 129S females for germline transmission of the targeted DISC1 allele.

#### Mouse breeding

Heterozygous DISC1-Boymaw mice were bred with wildtype mice to generate wildtype and heterozygous mice on 129S genetic background (129S2/SvPasCrl, Charles River) for all molecular studies. Different cohorts of mice were generated on the F1 hybrid (129S/C57) and 129S genetic backgrounds for behavioral studies. PCR was used for DISC1-Boymaw mouse genotyping (DISC1-Common-R: 5’ TAACAACAGCCAGTGTGCAAG 3’; DISC1-Neo-F: 5’ GGTGGGCTCTATGGCTTCTGA 3’; DISC1-F: 5’ TGCATTCACATGTGTTGCCTT TG 3’). Mice were housed in groups (2 to 4 mice per cage) in a climate-controlled animal colony with a reversed day/night cycle. Behavioral testing occurred during the dark cycle. Food (Harlan Teklab, Madison, WI) and water were available *ad libitum*, except during behavioral testing. All behavioral testing procedures were approved by the UCSD Institutional Animal Care and Use Committee (permit number: A3033–01) prior to the onset of the experiments. Mice were maintained in American Association for Accreditation of Laboratory Animal Care (AALAC) approved animal facilities. For behavioral studies, all mice were 2- to 6-month old siblings, and the age range of the birth date between all mice was less than a week.

#### Hematological analysis

A cohort of wildtype and heterozygous DISC1-Boymaw mice was generated on 129S background. Male mice were anesthetized with isofluorane before blood collection. Submandibular venous lancets (Goldenrod 5mm) were used to collect approximately 100uL into EDTA anticoagulated 0.5 mL Microtainer tubes (Becton-Dickinson). Hematological analysis was conducted using the Hemavet 950 FS at the UC San Diego Hematology Core Laboratory. 6 adult wildtype male mice and 6 heterozygous male DISC1-Boymaw siblings were first used for hematological analysis. To replicate the findings, another independent group of wildtype mice and their heterozygous DISC1-Boymaw siblings (N=4, for each genotype) was used.

#### RNA, DNA extraction and RT-PCR analysis

DNA and RNA were extracted from cells and mouse brains using Trizol method. RNA expression of the DB7 fusion gene was examined with RT-PCR using the primers: human DISC1 e1-f: 5’GCGGCGTGAGCCACCGCGCAG3’ human DISC1 e2-r: 5’GAACCGGAACAGTGTGCCCAC3’.

Mouse endogenous *disc1* RNA expression was examined for alternative splicing of exon 6 using primers:

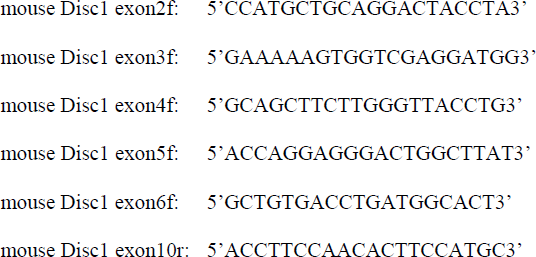

For the real-time RT-PCR quantification of mRNA transcripts of *Gad67*, *Nmdar1*, and *Psd95* genes, cDNA was first synthesized with random hexamers using the Superscript III First-Strand Synthesis Kit (Invitrogen). A comparative Ct method was used for quantification with SYBR Green, according to the manufacturer’s protocol (Bio-Rad CFX384). Beta-actin was used as a reference control for normalization. The following primers were used in Q-PCR:

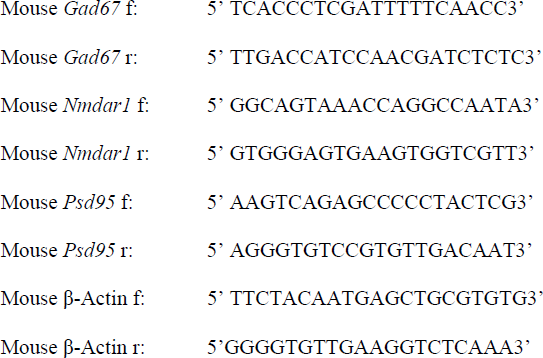

#### Immunohistochemical and Immunocytochemical staining and Western Blot

Immunohistochemical staining of parvalbumin, Gad67, Nmdar1 were conducted in mouse brain paraffin sections as described in our previous publication (83). For immunocytochemical staining, primary neurons were fixed in 3.7 % formalin/PBS at room temperature for 10 min. After 1 h of blocking with 5% goat serum containing 0.2% Triton X-100 at room temperature, the slides were incubated with mouse monoclonal anti-DISC1 (3D4) (1:1,000 dilution, gift from Dr. Carsten Korth) and rabbit polyclonal anti-MAP2 (1:2,000 dilution, PRB-547C, Covance) antibodies at 4 °C overnight. After rinsing three times with PBST, the slides were incubated with Alexa Fluor ^^®^^ 568 goat anti-mouse IgG (1:500 dilution, Invitrogen) and Alexa Fluor ^^®^^ 488 goat anti-rabbit IgG (1:500 dilution, Invitrogen) for 1 hr at room temperature. The slides were washed three times with PBST each for 5 min, and mounted with VECTASHIELD^®^ Hard Set™ Mounting Medium with DAPI (Vector, CA). The DB7 fusion proteins were detected in Western blot analysis of primary neuron culture using rabbit polyclonal anti-DISC1 antibody (1:500 dilution, ab55808, Abcam) recognizing N-terminal residues of human DISC1proteins. Expression of Gad67, Nmdar1, Psd95, synaptophysin, and β-actin in mouse brain and primary culture of neuronal cells was examined using the antibodies: mouse monoclonal anti-Gad67 (1:10,000 dilution, G5419, sigma); mouse monoclonal anti-Nmdar1 (1:5,000 dilution, 556308, BD Biosciences); mouse monoclonal anti-Psd95 (1:3,000 dilution, Ab99009, Abcam); rabbit polyclonal anti-synaptophysin (1:3,000 dilution, 5461S, Cell signaling); mouse monoclonal anti-β-actin (1:5,000 dilution, sc-47778, Santa Cruz Biotechnology).

#### SUnSET in mice

The *in vivo* SUnSET experiments were conducted as described (24). Sex-, and age-matched two-month old sibling wildtype and heterozygous mice were anaesthetized. Unilateral intracerebroventricular (ICV) injection of puromycin (25 ug/2.5ul) was performed with coordinates: -0.22 mm anterioposterior, -1.0 mm mediolateral, and -2.4 mm dorsoventral. Mice were sacrificed by cervical dislocation 1 hr after the injection. The hippocampus was dissected for total protein extraction. Equal amounts of protein (70 μg) were loaded on PAGE gel for Western blot analysis. Anti-puromycin antibody (1:1,000)(12D10, Millipore) was used to detect incorporation of puromycin in newly synthesized proteins. Blots were then stripped and re-probed with ant-β-actin antibody. Image J was used to quantify the incorporation of puromycin.

#### RNA in situ hybridization

We developed a modified protocol for chromogenic RNA *in situ* hybridization according to branched DNA amplification assays (84, 85). Oligonucleotide adaptors were added to the 5’ end of three 18S rRNA probes (5’TCACTGTACC GGCCGTGCGT3’; 5’CCTAGCTGCGGTATCCAGGC3’; 5’GCGATCGGCCCGAGGTTATC3’). In brief, paraffin sections were dewaxed, rehydrated, and partially digested with pepsin (4 mg/ml) at 37 °C for 10 min. After PBS washing, sections were pre-hybridized with Quick-hyb Solution (Stratagene, 201220) with 0.1M DTT, and 50 µg/ml denatured ssDNA. Pre-hybridization was conducted at 40 °C for 15 min in pre-heated wet plastic chamber. After pre-hybridization, probes were added to the final concentration of 200 ng/ml. Hybridization was conducted at 40 °C overnight in pre-heated wet pyrex chamber. After hybridization, sections were washed, and hybridized with oligonucleotide amplifier labeled with biotin. Horseradish Peroxidase-conjugated avidin (Thermo Scientific, 21123) was used to bind biotin at final concentration of 5.5 ug/ml. Chromogenic reaction was performed using DAB Peroxidase Substrate Kit (Catalog# SK-4100, Vector Labs).

#### Acoustic startle

Startle reactivity and prepulse inhibition (PPI) were measured with startle chambers (SR-LAB, San Diego Instruments, San Diego, CA) using 65 dB background noise, three types of prepulse-pulse trials (69 dB, 73 dB, 81dB) and pulse-alone (120 dB) (86). The session started with 5 pulse-alone trials at 120 dB intensity to stabilize startle. The prepulse testing block contained 3 prepulse trial types in which the 69 dB prepulse onset preceded the 120 dB pulse by 25, 50, 100, 200, and 500 msec. Prepulse trials and pulse alone trials were presented in a pseudorandom order. The session ended with 5 pulse-alone trials. There was an average of 15 s (range: 12–30 s) between trials. Mice were assigned in a pseudo-random order, and placed in the startle chambers, where a 65-dB background noise level was presented for a 10-min acclimation period and continued throughout the test session.

#### Tail suspension

The tail suspension test was conducted as described previously with some modifications (45, 46). In brief, mice were suspended by the tail with adhesive tape placed 1–2 cm from the end of the tail. The mice were visually isolated at the center of a white box. Mice were suspended for 6 min, and the duration of immobility - during which the mice did not produce any apparent voluntary movements - was recorded by an observer blind to genotype.

#### Saccharin preference

The saccharin preference test was conducted as described with some modifications (87). In brief, mice were separated and housed, 2 or 3 per cage, by genotype. Saccharin sodium salt hydrate (Sigma) was dissolved in tap water to 1.0%. Each cage was supplied with one bottle of saccharin solution and one bottle of normal drinking water. The bottles were weighed before placement in each cage and 24, 48, 72, and 96 hr thereafter. Preference for the saccharin solution was calculated as the percentage of saccharin solution consumed out of the total liquid consumption each day. Food was available *ad-libitum* during the course of the experiment.

#### Behavioral pattern monitor

Spontaneous behavioral data were recorded using the mouse BPM (San Diego Instruments, San Diego, CA), as described previously (44, 88). In brief, a single chamber consists of a 30.5× 61× 38-cm area, with a Plexiglas hole board floor that was equipped with floor holes in the front, middle, and rear parts of the floor and eight wall holes (three along each side of the long walls and two holes in the front and back walls). Mice were tested during the dark phase of their light cycle. During testing, a white noise generator produced background noise at 65 dB. The measurement of transitions, center time, and spatial coefficient of variation were based on the nine divided regions of the chambers. The status of the photobeams was sampled every 0.1 sec. The session lasted 60 min. Raw data were transformed into the location of the animal (in X–Y coordinates), whether holepoking or rearing occurred (events), and the duration of each event (time). The chambers were cleaned thoroughly between testing sessions.

#### Ketamine treatment

Ketamine was dissolved in saline, and administered i.p. at a volume of 5 ml/kg with a dosage of 60 mg/kg immediately prior to the start of the BPM session. A within-subjects design was used for the ketamine studies. After administration of ketamine, mouse locomotor activities were recorded for 60 min in BPM chambers.

#### Statistical analysis

All data were first tested for normal distribution using the Kolmogorov-Smirnov test before calculation of differences. For statistical analyses, repeated measures analysis of variance (ANOVA) with genotypes as a between-subjects factor and drug treatment, block, and prepulse intensity as within-subjects factors were performed on the %PPI data and locomotor activity (e.g., total distance). *Post hoc* analyses were carried out using Newman-Keuls or Tukey’s test. Alpha level was set to 0.05. All statistical analyses were carried out using the BMDP statistical software (Statistical Solutions Inc., Saugus, MA).

## Acknowledgments

We thank Drs. Carsten Korth and Nick Brandon for generously providing anti-DISC1 antibodies.

## Contributions

X.Z. conceived, performed, analyzed, and wrote the studies. B.J., K.H., M.K., L.Z., and J.Y. performed and analyzed the research. M.G. provided critical discussion and comments on the studies.

